# Expansion Revealing: Decrowding Proteins to Unmask Invisible Brain Nanostructures

**DOI:** 10.1101/2020.08.29.273540

**Authors:** Deblina Sarkar, Jinyoung Kang, Asmamaw T Wassie, Margaret E. Schroeder, Zhuyu Peng, Tyler B. Tarr, Ai-Hui Tang, Emily Niederst, Jennie. Z. Young, Li-Huei Tsai, Thomas A. Blanpied, Edward S. Boyden

**Author notes:** These authors contributed equally (Deblina Sarkar, Jinyoung Kang, Asmamaw T Wassie).

## Abstract

Cells and tissues are made out of nanoscale building blocks, such as proteins, organized in crowded nanostructures. We show that many biomolecules, and the nanostructures in which they are embedded, may be invisible to prior imaging techniques, due to the inaccessibility of labels (e.g., antibodies) to biomolecules embedded within such crowded structures. We developed a technology, expansion revealing (ExR), which isotropically decrowds proteins from each other, to enable their labeling. We use ExR to discover the alignment of presynaptic calcium channels with postsynaptic machinery in intact brain circuits, as well as the existence of periodic amyloid nanoclusters containing ion channel proteins in Alzheimer’s model mice. Thus, ExR reveals nanostructures within complex biological specimens that were not previously visualizable, and may find broad use in biology.

**One Sentence Summary:** De-crowding biomolecules with Expansion Revealing unmasks fundamentally new nanostructures in intact brain tissue, which remained invisible otherwise.

## Main Text

Understanding in 3D how proteins are configured into structures within cells and tissues, with nanoscale precision, is key to understanding biology. Super-resolution microscopy has offered the capability of combining biomolecular recognition, through staining with target protein-specific antibodies, with nanoscale optical resolution (*1*). However, such techniques are limited by the accessibility of labels to the biomolecules to be imaged, because it is the labels that are visualized, and not the biomolecules themselves. Many crowded, biomolecule-rich structures (such as synapses) are at the core of biological functionality and disease states, with inter-protein distances smaller than the size of antibodies, and which thus may prohibit access to biomolecules of interest by labels in such crowded environments. Overcoming this issue requires a technology which can not only achieve super-resolution down to the low tens of nanometers, but also allow decrowding of biomolecules. Could such a technology reveal fundamentally new biological nanostructures, not previously visible?

Here we develop and validate such a technology, and systematically answer this question. We reasoned that in order to decrowd proteins from each other, we could leverage the spatial expansion property of expansion microscopy (*2, 3*) (in which cells and tissues are densely permeated by an even mesh of swellable hydrogel, and then expanded, to obtain enhanced resolution on ordinary microscopes). While many papers have alluded to decrowding (*4–8*), driven by the appeal of the possibility of better access to crowded proteins by labels, the key problem is that decrowding involves a sort of inherent contradiction: if expanding proteins away from each other reveals new structures, how does one know if the new structures are real? Given that previous claims about decrowding have not been substantiated by data showing this, we here set out to create a technology that achieves decrowding, and to show that it is trustworthy and useful. To establish decrowding and at the same time achieve super-resolution on par with the best classical super-resolution techniques (down to ~20 nm resolution), we combined the expansion principle with a novel scheme for biomolecular preservation, to build a technology that we here call expansion revealing (ExR). By direct comparison of biomolecular structures where labeling was done in the normal, crowded environment vs. after the decrowding effect of ExR, with both being imaged at the same level of resolution (a comparison uniquely enabled by the new ExR methodology), we demonstrate that ExR indeed is revealing new biological information, and indeed, can lead to the discovery of fundamentally new nanostructures in intact brain tissue, which would otherwise have remained invisible. Importantly, through careful comparisons of pre- vs. post-expansion staining enabled by ExR, we show that the newly discovered nanostructures reflect biologically meaningful findings, as opposed to artifacts of decrowding chemistry. Thus we anticipate ExR will enable the visualization of a variety of previously undescribed biological nanostructures; we illustrate the capabilities of ExR by unmasking key components of synaptic nanocolumns as well as periodic nanostructures potentially of relevance to Alzheimer’s disease.

### Expansion revealing: a technology that enables simultaneous super-resolution and decrowding

To achieve super-resolution down to the low tens of nanometers, two rounds of expansion (or 15-20x expansion factor) is required. One would ideally minimally alter the proteins during the expansion process, while decrowding them from one another as much as possible. To satisfy these conflicting requirements, ExR (Fig. 1A-C) achieves this with a novel scheme. We reasoned that one could expand a brain specimen through one round of gelation and expansion, and that this first swellable hydrogel could then be further expanded if we formed a second swellable hydrogel in the space opened up by the first expansion, and then swelled the specimen a second time (see Methods for details). The proteins, being anchored to the first hydrogel throughout the entire process, would be retained because the original hydrogel would be further expanded by the second swellable hydrogel, and not cleaved or discarded. This strategy avoids the need to transfer biomolecular information from the first hydrogel to the second one, which is challenging and undesirable, since it requires either DNA intermediates as in iterative Expansion Microscopy (iExM) (*9*) (which necessitates the loss of original biomolecules) or second fixation as in a recent iExM variant (*6*) (which has not been validated in terms of protein decrowding). Thus, ExR imposes no extra processing steps on the proteins than is required for the first expansion. In the final step, the proteins can be antibody labeled (Fig. 1C), after decrowding and before imaging, enabling visualization of proteins that would have been missed if stained in the crowded state (Fig. 1B).

**Fig. 1.**
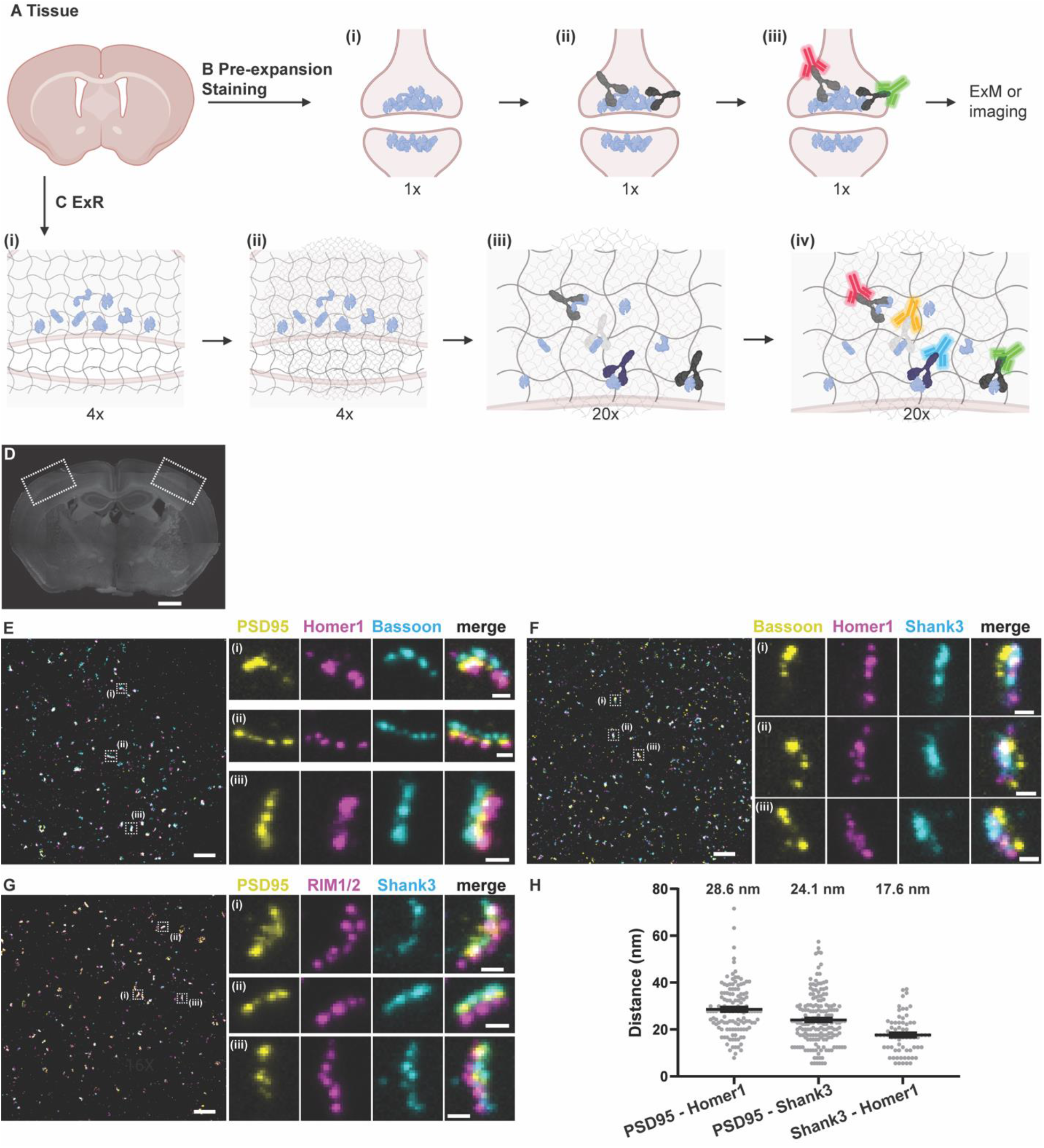
Expansion Revealing (ExR), a technology for decrowding of proteins through isotropic protein separation. (A) Coronal section of mouse brain before staining or expansion. (B) Conventional antibody staining may not detect crowded biomolecules, shown here in pre- and post-synaptic terminals of cortical neurons. (Bi) Crowded biomolecules before antibody staining. (Bii) Primary antibody (Y-shaped proteins) staining in non-expanded tissue. Antibodies cannot access interior biomolecules, or masked epitopes of exterior biomolecules. (Biii) Secondary antibody (fluorescent green and red Y-shaped proteins) staining in non-expanded tissue. After staining, tissue can be imaged or expanded using earlier ExM protocols, but inaccessible biomolecules will not be detected. (C) Post-expansion antibody staining with ExR. (Ci) Anchoring and first gelation step. Specimens are labeled with gel-anchoring reagents to retain endogenous proteins, with acrylamide included during fixation to serve as a polymer-incorporatable anchor, as in refs (*5, 7*). Subsequently, the specimen is embedded in a swellable hydrogel that permeates densely throughout the sample (gray wavy lines), mechanically homogenized via detergent treatment and heat treatment, and expanded in water. (Cii) Re-embedding and second swellable gel formation gelation. The fully expanded first gel (expanded 4x in linear extent) is re-embedded in a charge-neutral gel (not shown), followed by the formation of a second swellable hydrogel (light gray wavy lines). (Ciii) 20x expansion and primary antibody staining. The specimen is expanded by another factor of 4x, for a total expansion factor of ~20x, via the addition of water, then incubated with conventional primary antibodies. Because expansion has decrowded the biomolecules, conventional antibodies can now access interior biomolecules and additional epitopes of exterior molecules. (Civ) Post-expansion staining with conventional fluorescent secondary antibodies (fluorescent blue and yellow Y-shaped proteins, in addition to the aforementioned red and green ones) to visualize decrowded biomolecules. Schematic created with BioRender.com. (D) Low-magnification widefield image of a mouse brain slice stained with DAPI, with box highlighting the somatosensory cortex used in subsequent figures for synapse staining (Scale bar, 500 μm), and (E-G) confocal images (max intensity projections) of representative fields of view (cortical L2/3) and specific synapses after ExR expansion and subsequent immunostaining using antibodies against (E) PSD95, Homer1, Bassoon; (F) Bassoon, Homer1, Shank3; and (G) PSD95, RIM1/2, Shank3. Scale bar, 1 μm, left image; 100 nm, right images; in biological units, i.e. the physical size divided by the expansion factor, throughout the paper unless otherwise indicated. (H) Measured distance between centroids of protein densities of PSD95 and Homer1, PSD95 and Shank3, and Shank3 and Homer1, in synapses such as those in panels e-g. The mean distance (again, in biological units) between PSD95 and Homer1 is 28.6 nm (n = 126 synapses from 3 slices from 1 mouse), between PSD95 and Shank3 is 24.1 nm (n=172 synapses from 3 slices from 1 mouse), and between Shank3 and Homer1 is 17.6 nm (n=70 synapses from 3 slices from 1 mouse). Plotted is mean +/- standard error; individual grey dots represent the measured distance for individual synapses.

We first quantified the global isotropy of the expansion process in ExR, and found a similar low distortion (i.e., of a few percent over length scales of tens of microns, Fig. S1) as we did for previous ExM protocols (*3, 4, 9–11*). As with previous protocols, to measure effective resolution, we focused on synapses, given both their importance for neural communication and utility as a super-resolution testbed, staining post-expansion cortical synapses (Fig. 1D) with antibodies against the pre-synaptic protein Bassoon and the post-synaptic proteins PSD95 and Homer1 (Fig. 1E-G). We found that when we measured the mean distance between domains containing the proteins PSD95, Homer1, and Shank3, we obtained values (Fig. 1H) that were very similar to classical results found using STORM microscopy(*12*) (note that our study focused on Shank3 and this earlier study focused on Shank1). Thus, ExR exhibits effective resolution, on the order of ~20 nm, comparable to our earlier iExM protocol, which did not retain proteins (*9*).

### High-fidelity enhancement of synaptic protein visualization via ExR

To gauge whether ExR could reveal nanostructures in the brain that were not visible without decrowding, and to further probe whether it incurred any costs in terms of decreased resolution or added distortion relative to classical staining (i.e. pre-expansion staining and thus, no decrowding), we devised a strategy where we would stain brain slices pre-expansion with an antibody against a synaptic protein, in a fashion so that antibodies would be anchored to the expansion hydrogel for later visualization, then perform ExR, and then restain the same proteins with the same antibody a second time, thus enabling a within-sample comparison of a given protein across both conditions (with and without decrowding) at the same level of resolution, which could reveal any nanostructural differences. For pre-expansion staining (see Methods for details), we immunostained mouse brain slices containing somatosensory cortex (Fig. 2A) with primary antibodies followed by 6-((acryloyl)amino)hexanoic acid, succinimidyl ester (abbreviated AcX)-conjugated secondary antibodies, so that these antibodies could be attached to the swellable hydrogel for post-expansion tertiary antibody staining and visualization. This allowed us to compare pre-ExR staining to post-ExR staining, at the same resolution level, and for the same field of view, important for noticing any changes in nanostructural detail. We noted that Homer1 and Shank3 exhibited very similar visual appearances when we compared pre- vs. post-ExR staining (quantified below), so we designated these two stains as “reference channels,” that is, co-stains that could help define synapses for the purposes of technological comparison, and that we could use to help us gauge whether other proteins were becoming more visible at synapses.

**Fig. 2.**
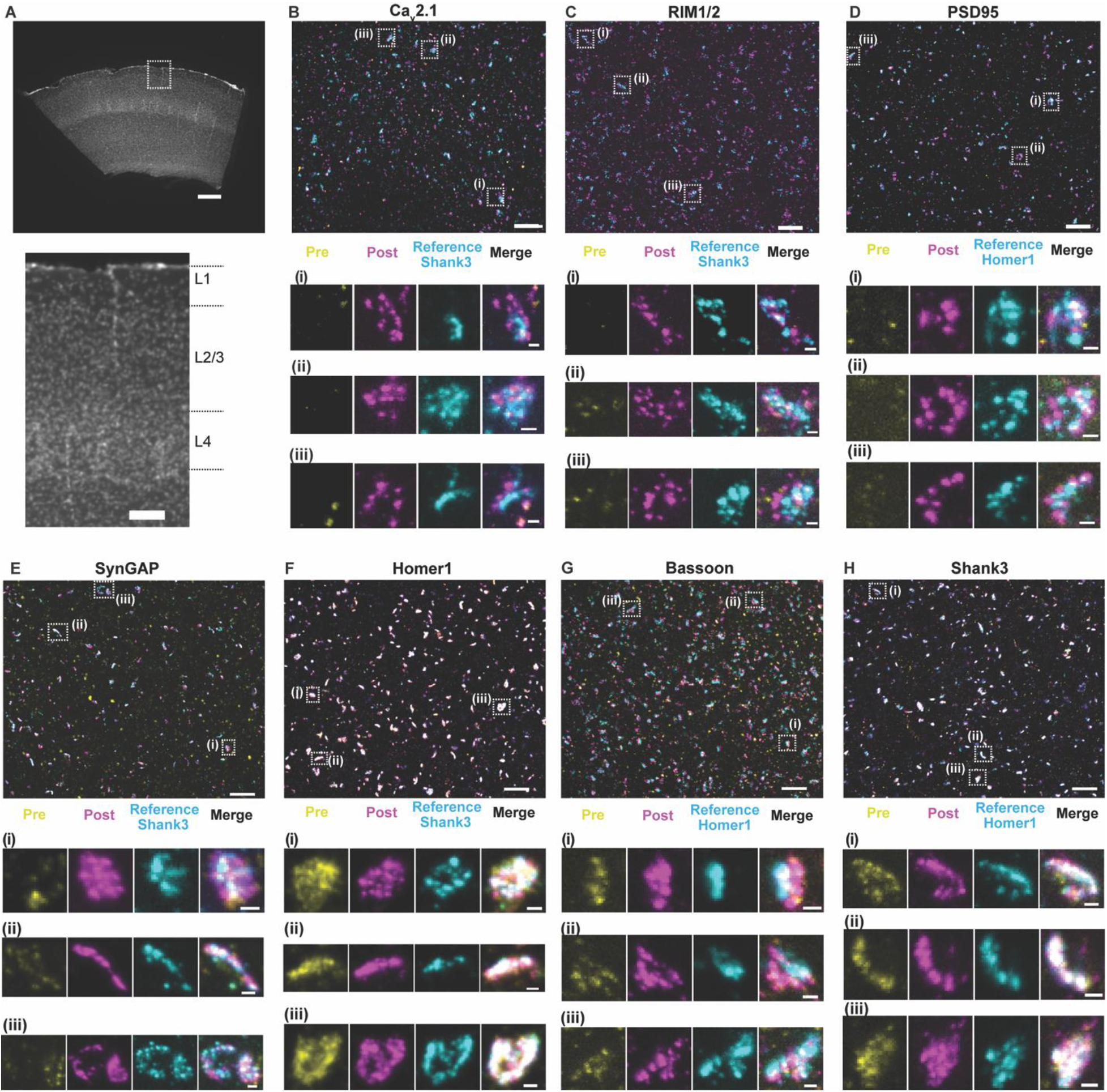
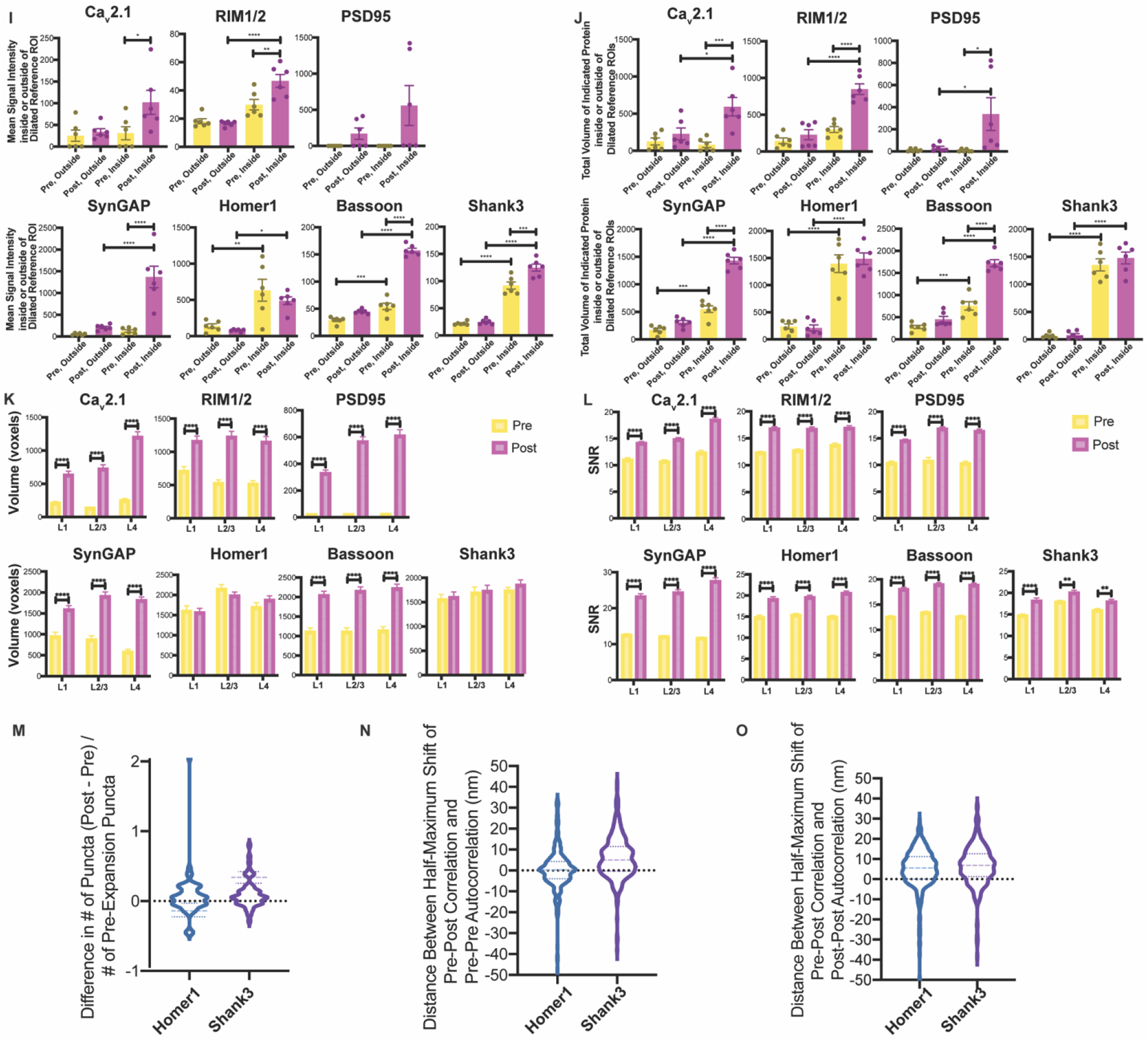
Validation of ExR enhancement and effective resolution in synapses of mouse cortex. **(A)** Low-magnification widefield image of a mouse brain slice with DAPI staining showing somatosensory cortex (top) and zoomed-in image (bottom) of boxed region containing L1, L2/3, and L4, which are imaged and analyzed after expansion further in panels **B**-**H**. (Scale bar, 300 μm (top) and 100 μm (bottom)). **(B-H)** Confocal images of (max intensity projections) of specimen after immunostaining with antibodies against Cav2.1 (Ca^2+^ channel P/Q-type) **(B)**, RIM1/2 **(C)**, PSD95 **(D)**, SynGAP **(E)**, Homer1 **(F)**, Bassoon **(G)**, and Shank3 **(H),** in somatosensory cortex L2/3. For pre-expansion staining, primary and secondary antibodies were stained before expansion, the stained secondary antibodies anchored to the gel, and finally fluorescent tertiary antibodies applied after expansion to enable visualization of pre-expansion staining. For post-expansion staining, the same primary and secondary antibodies were applied after ExR. Antibodies against Shank3 **(B, C, E, F)** or Homer1 **(D, G, H)** were applied post-expansion as a reference channel. Confocal images of cortex L2/3 (top) show merged images of pre- and post-expansion staining, and the reference channel. Zoomed-in images of three regions boxed in the top image (**i-iii,** bottom) show separate channels of pre-expansion staining (yellow), post-expansion staining (magenta), reference staining (cyan), and merged channel. (Scale bar, 1.5 μm (upper panel); 150 nm (bottom panel of i-iii).) **(I, J)** Quantification of decrowding in a set of manually identified synapses. Statistical significance was determined using Sidak’s multiple comparisons test following ANOVA (*P≤0.05, **P≤0.01, ***P < 0.001, ****P < 0.0001, here and throughout the paper, and plotted is the mean, with error bars representing standard error of the mean (SEM), here and throughout the paper). (**I)** Mean signal intensity inside and outside of (i.e., nearby to) dilated reference ROIs for pre- and post-expansion stained manually-identified synapses (**Table S1** for numbers of technical and biological replicates)). Data points represent the mean across all synapses from a single field of view. (**J)** Total volume (in voxels; 1 voxel = 17.16×17.15×40, or 11,779, nm^3^) of signals inside and outside of dilated reference ROIs, in both cases within cropped images containing one visually identified synapse (**Fig. S3h**), for pre- and post-expansion stained manually-identified synapses (**Table S1** for numbers of biological and technical replicates). Data points represent the mean across all synapses from a single field of view. **(K)** Mean voxel size and **(L)** mean signal-to-noise (SNR) ratio of pre- and post-expansion immunostaining showing 7 proteins in somatosensory cortex regions L1, L2/3, and L4 of 2 mice. Plotted is mean and SEM. To compare the 3D voxel size and SNR of pre- and post-expansion stained synapses for each of the seven proteins, three t-tests (one for each layer) were run (n = 49-70 puncta per layer from 2 mice; **Table S1** for replicate numbers). Statistical significance was determined using multiple t-tests corrected using the Holm-Sidak method, with alpha = 0.05. **(M)** Population distribution (violin plot of density, with a dashed line at the median and dotted lines at the quartiles) of the difference in the number of synaptic puncta between post- and pre-expansion staining channels for Homer1 and Shank3 (Homer1, n = 304 synapses from 2 mice; Shank3, n = 309 synapses from 2 mice. **(N)** Population distribution of the difference in distance (in nm) between the half-maximal shift for pre-pre autocorrelation and post-pre correlation (calculated pixel-wise between intensity values normalized to the minimum and maximum of the image, see Methods) averaged over x-, y-, and z-directions (x- and y-directions being transverse, z-direction being axial) for Homer1 and Shank3 (Homer1, n = 303 synapses from 2 mice, Shank3, n = 305 synapses from 2 mice). **(O)** Population distribution of the difference in distance (in nm) between the half-maximal shift for post-post autocorrelation and post-pre correlation (calculated pixel-wise between intensity values normalized to the minimum and maximum of the image) averaged over x-, y-, and z-directions for Homer1 and Shank3 (Homer1, n = 303 synapses from 2 mice, Shank3, n = 305 synapses from 2 mice).

We chose 7 synaptic proteins important for neural architecture and transmission for this experiment – the presynaptic proteins Bassoon, RIM1/2, and the P/Q-type Calcium channel Cav2.1 alpha 1A subunit, and the postsynaptic proteins Homer1, Shank3, SynGAP, and PSD95 (Fig. 2B-H), staining for each protein along with a reference channel stain (note, when we imaged Homer1, we used Shank3 as the reference channel, and vice versa). All seven proteins exhibited well-defined images when post-expansion stained, with the geometry reflecting characteristic synaptic shapes. For example, a presynaptic and a postsynaptic stain (Fig. 2B, 2C, 2G) revealed parallel regions with a putative synaptic cleft in between, although note that this was only visible if the synapse was being imaged from the side; if a synapse was being imaged with the axial direction of the microscope perpendicular to the synaptic cleft, then the image may look more disc-shaped (*9, 12, 13*). In many cases, however, post-ExR staining revealed more detailed structures of synapses compared to what was visualized with pre-expansion staining – for example, calcium channels (Fig. 2B), RIM1/2 (Fig. 2C), PSD95 (Fig. 2D), SynGAP (Fig. 2E), and Bassoon (Fig. 2G) appeared more prominent post-expansion than pre-expansion. Applying a conventional antigen retrieval protocol did not result in such improvements (Fig. S2), suggesting that the decrowding effect observed via ExR was indeed due to expansion, and not simply due to denaturation or antigen retrieval-like effects associated with other aspects of the ExR process.

To quantitate the improvement in staining enabled by ExR, we measured the amplitude and volume of each synaptic protein stain, both within, and just outside of, identified synapses. First, we manually identified between 49-70 synapses (Table S1 for exact numbers) per ~350×350×20 μm3 (in physical units, e.g. what is actually seen through the microscope lens) field of view, choosing the largest and brightest synapses based on reference channel staining (i.e., Homer1 or Shank3). We developed an automated method to segment synaptic puncta from nearby background. Briefly, we created binary image stacks for each channel using a threshold equal to a multiple of the average measured standard deviation of five manually-identified background regions (not blinded to condition), filtered 3D connected components based on size, and used dilated reference channel ROIs to segment putative synaptic puncta (Fig. S3 and Methods). We dilated reference channel-defined ROIs to relax the requirement of exact colocalization of pre-synaptic proteins with a post-synaptic reference. We found that post-expansion staining increased signal intensity and mean total volume of signal within dilated reference-channel defined ROIs, without meaningfully affecting background (i.e., signal just outside the dilated reference channel-defined ROIs) staining (Fig. 2I-J). In summary, all proteins except Homer1 and PSD95 showed significantly increased signal intensity of post-expansion staining within dilated reference ROIs, with minimal increase in the background (Fig. 2I; Table S2 for full statistical analysis). (We note that there was no change in signal intensity in the background for PSD95, but a bimodal distribution of signal intensity increases in the foreground, potentially due to poor signal quality in one animal.) Similarly, all proteins exhibited increased total volume of post-expansion staining signals within dilated references ROIs, and minimal or no increase in the volume of background ROIs (Fig. 2J; Table S3 for full statistical analysis). Because Homer1 and Shank3 had the smallest changes in pre- vs. post-expansion staining, they were chosen as reference channels to indicate the locations of putative synapses for pre- vs. post-expansion test stain comparison, as mentioned above. Of course, different antibodies may bind to different sites on a target protein, and we found that a different antibody against PSD95 (from Cell Signaling Technology, product number CST3450S) than the one used in Figure 2 (Thermofisher, product number MA1-046) showed similar signal intensity and volume when compared pre- vs. post-staining (Fig. S4); perhaps, in future studies, the use of multiple antibodies against different parts of a single protein, in pre- vs. post-expansion comparisons, could be used to gauge the density of the environment around different parts of that protein.

We further analyzed synaptic protein signals pre- vs. post-expansion in the context of different cortical layers. We quantified volume and signal-to-noise ratio (SNR; signal intensity divided by standard deviation of the background) of each protein in 3D synaptic structures by binarizing signals over a threshold (a multiple of the standard deviation of the intensity within a manually-selected background region; see Methods) and selecting putative synapses in each ~350×350×20 μm3 field of view imaged above, comparing the values of pre- vs. post-expansion staining in each layer of somatosensory cortex (L1, L2/3 and L4, respectively) (Fig. 2K-L). Post-expansion volumes exhibited larger volumes in each layer for all of the stains except for the reference channel stains, and improved SNR for all synaptic proteins (Tables S4 and S5 for full statistics).

In previous work, we showed using the iterative expansion microscopy (iExM) protocol, which uses pre-expansion staining, that we could achieve effective resolutions of ~20 nm, with low distortion due to the expansion process (*9*). As another independent way to gauge the potential distortion obtained by staining with antibodies post-expansion, we compared the shapes of synaptic puncta as seen with the pre-expansion stain, with the shapes as seen with the new post-expansion stain, using the within-sample dual staining method used in Figure 2a-l. We compared various properties of synaptic puncta between pre- and post-expansion staining conditions, using the reference proteins Homer1 and Shank3 since they had similar intensities and volumes when comparing pre- and post-expansion datasets, and therefore might be appropriate for comparing shape features across these conditions (Fig. S5). In summary, we did not see significant distortion being introduced by post-expansion staining; for example, we found no significant difference in the number of synaptic puncta when we compared pre- vs. post-expansion staining in the same sample (Fig. 2M), and when we measured the shift in puncta positions between pre- and post-stained conditions, we observed average shifts of <10nm (in biological units, i.e. physical size divided by the expansion factor) between post- and pre-expansion staining for both Homer1 and Shank3 (Fig. 2N-O; Table S4 for full statistics). Thus ExR preserves, relative to classical pre-expansion staining, the locations of proteins with high fidelity for the purposes of post-expansion staining.

### Synaptic nanocolumns coordinated with calcium channel distributions

Coordinating pre- and post-synaptic protein arrangement in a nanocolumn structure which aligns molecules within the two neurons contributes to precision signaling from presynaptic release sites to postsynaptic receptor locations(*13, 14*), as well as to the long-term plasticity of synaptic function (*15*). Given that ExR is capable of unmasking synaptic proteins that are otherwise not detectable, with nanoscale precision, we next sought to explore the nanocolumnar architecture of pre- and postsynaptic proteins, with a focus on important molecules that have not yet been explored in this transsynaptic alignment context. As noted above, ExR greatly helps with visualization of calcium channels, which of course are amongst the most important molecules governing the activation of synaptic release machinery, with nanometer-scale signaling contributing to the precision of synaptic vesicle fusion. However, the nanoscale mapping of calcium channels in the context of nanocolumnar alignment in intact brain circuits remains difficult (*16–18*) (Fig. 2B). We thus applied ExR to investigate whether calcium channels occupy nanocolumns with other pre- and post-synaptic proteins, such as the critical pre- and post-synaptic proteins RIM1/2 and PSD95, respectively, across the layers of the cortex (Fig. 3A-D). We first performed a 3D autocorrelation function (ga(r))-based test which provides information about the intensity distribution within a defined structure (see Methods). Any heterogeneity in the intensity distribution within the cluster will result in a ga(r)>1, and the distance at which the ga(r) crosses 1 can be used to estimate the size of the internal heterogeneity, here termed a nanodomain (*13*). In all cortical layers, our auto-correlation analysis shows that all 3 proteins explored exhibited a non-uniform arrangement forming nanodomains with average diameters of about 60-70nm (in biological units; Fig. 3E-H). To analyze the spatial relationship between the two distributions and the average molecular density of RIM1/2, PSD95 and Cav2.1 relative to each other, we performed a protein enrichment analysis, which is a measure of volume-averaged intensity of one channel as a function of distance from the peak intensity of another channel (see Methods). To more easily compare the extent of the enrichment between these proteins in each layer, we also calculated the enrichment index, which is an average of all enrichment values within 60 nm (biological units) of the peak of a designated channel. Our analysis shows that the centers of nanodomains of RIM1/2, PSD95 and Cav2.1 are enriched with respect to each other (Fig. 3I-P). Of particular interest, the nanoscale colocalization of Cav2.1 with RIM1/2 (and thus the vesicle site) likely minimizes the distance between the channels and the molecular Ca sensors that trigger vesicle fusion (Fig. 3Q-R), consistent with the physiological concept of nanodomain coupling that tunes the efficacy and frequency-dependence of neurotransmission (*19*). Furthermore, the precise alignment between RIM1/2 and PSD95 may reduce the distance that released neurotransmitter needs to diffuse before reaching postsynaptic receptors (Fig. 3R). Thus, these nanoscale arrangements may help to optimize the speed, strength, and plasticity of synaptic transmission. To the best of our knowledge, this is the first report of the 3D nano-architecture of voltage gated calcium channel distributions within the transsynaptic framework probed in intact brain circuitry.

**Fig. 3.**
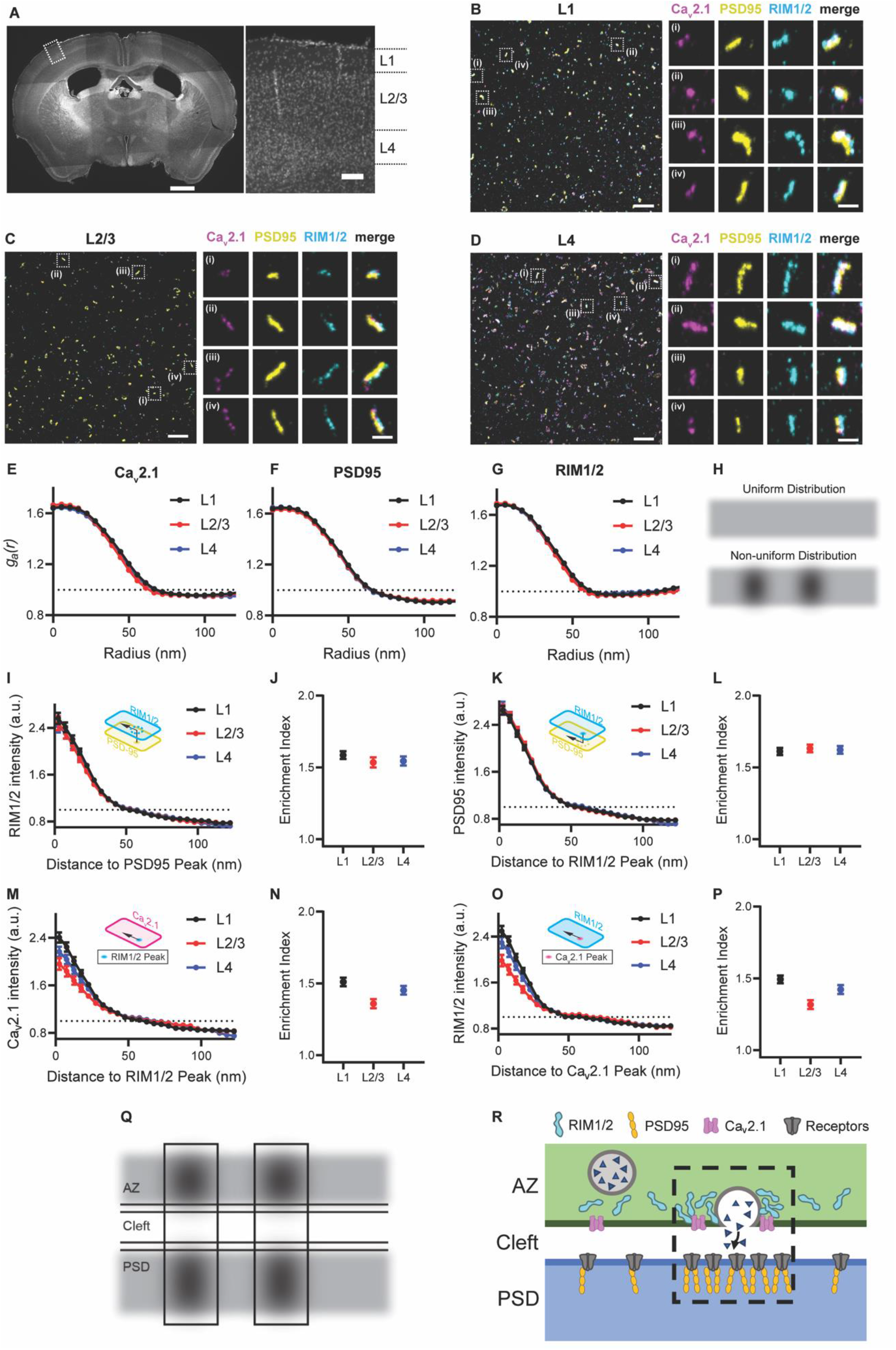
ExR reveals how calcium channel distributions participate in transsynaptic nanoarchitecture. (A) Low-magnification widefield image of DAPI stained mouse brain slice (left) and zoomed in view (right) of the boxed region showing layers 1-4 of the cortex. (Scale bar = 1000 μm (left panel of whole brain) and 100 μm (right panel of the layers 1-4). (B-D) show confocal images (max intensity projections) of layers 1, 2/3 and 4 respectively, after performing ExR and immunostaining with antibodies against Cav2.1 (calcium channel) (magenta), PSD95 (yellow) and RIM1/2 (cyan). In each of (B), (C) and (D), low magnification images are shown on the left while zoomed-in images of four regions, i-iv are shown on the right with separated channels for each antibody along with the merged image. (Scale bar = 1 μm (left panel) and 100 nm (right panels labeled i-iv)). (E-G) show the autocorrelation analysis for Cav2.1, PSD95 and RIM1/2 respectively for different layers, from which it is evident that these proteins have non-uniform distribution forming nanoclusters instead of having a uniform distribution as illustrated in (H). (I), (K), (M) and (O) show the enrichment analysis that calculates the average molecular density for RIM1/2 to the PSD95 peak, PSD95 to the RIM1/2 peak, Cav2.1 to the RIM1/2 peak and RIM1/2 to the Cav2.1 peak respectively while (J), (L), (N) and (P) show the corresponding mean enrichment indexes respectively. Error bars indicate SEM. (q) Schematic illustration represents that the nanoclusters of any two proteins (RIM1/2, PSD95 and Cav2.1) are aligned with nanoscale precision with each other. This may lead to efficient use of calcium ions in vesicle fusion as calcium channels are located close to vesicle fusion sites (dictated by RIM1/2) as well as efficient transfer of neurotransmitters across the synapse as the vesicle fusion site is located directly opposite to the location of receptor sites (PSD95 being a scaffolding protein) as schematically illustrated in (R).

### Periodic amyloid nanostructures in Alzheimer’s model mouse brain

In addition to densely crowded proteins in healthy functioning compartments like synapses, densely crowded proteins appear in pathological states like Alzheimer’s disease. Protein aggregates known as β-amyloid are thought to play roles in synaptic dysfunction, neurodegeneration and neuroinflammation (*20*). However, the densely packed nature of these aggregates may make the nanoscale analysis of their ultrastructure within intact brain contexts difficult to understand. To understand the nanoarchitecture of β-amyloid in the cellular context of intact brain circuitry, we applied ExR to the brains of 5xFAD Alzheimer’s model mice (Fig. 4A), as this widely used animal model of Alzheimer’s exhibits an aggressive amyloid pathology (*21*). We employed two different commercially available antibodies for β-amyloid, 6E10 (which binds to amino acid residues 1-16 of human Aβ peptides) and 12F4 (which is reactive to the C-terminus of human Aβ and has specificity towards Aβ42). We were particularly interested in investigating the relationship of amyloid deposits and white matter tracts, as these regions have been implicated in human imaging data (*22–25*), but are less investigated in mouse models. We previously reported the accumulation of Aβ aggregates along the fornix, the major white matter tract connecting the subiculum and mammillary body, early in disease progression (*25*). We co-stained the amyloid antibodies along with the axonal marker SMI-312, and compared pre-expansion staining with that obtained after ExR. Plaques appeared to be larger, and to have finer-scaled features in post-expansion staining, than in pre-expansion staining (Fig. 4B–4C), when visualized by either 6E10 or 12F4 antibodies, and thus the post-expansion staining may unveil aspects of plaque geometry that are not easily visualized through traditional means. Additionally, post-ExR staining revealed detailed nanoclusters of β-amyloid that were not seen when staining was done pre-expansion (Fig. 4B-C). These nanodomains of β-amyloid appeared to occur in periodic structures (Fig. 4B-C, subpanels (i)-(iv)). Co-staining with 2 different β-amyloid antibodies, D54D2 (which binds to isoforms Aβ37, Aβ38, Aβ40, and Aβ42) along with 6E10 or 12F4 in ExR-processed 5xFAD fornix showed similar patterns of periodic nanostructures, which means the observed periodic nanostructures are likely not composed of specific isoforms (Fig. S6). These nanodomains, and periodic structures thereof, were not visualized through pre-expansion staining, which was confirmed using unexpanded tissue as well (Fig. S7). To see whether the periodic nanostructures of β-amyloid revealed through ExR were just nonspecific staining artifacts of ExR, we performed ExR on wild type (WT) mice as a control, which should not have any labeling for human β-amyloid. As is clear from Figure 4D, no β-amyloid structures were observed in ExR-processed WT brain slice. Our quantitative analysis examining the volume of amyloid in ExR-processed WT and 5xFAD mice confirms that there is indeed a large amount of amyloid volume occupied in 5xFAD samples, but essentially no such volume occupied in ExR-processed WT mice (Fig. 4E). The lack of amyloid staining in ExR-processed WT mice makes it unlikely that the staining seen in ExR-processed 5xFAD mouse brain is nonspecific, and thus it is likely that the effect of ExR was to decrowd dense amyloid nanostructures that ordinarily hamper antibody staining.

**Fig. 4.**
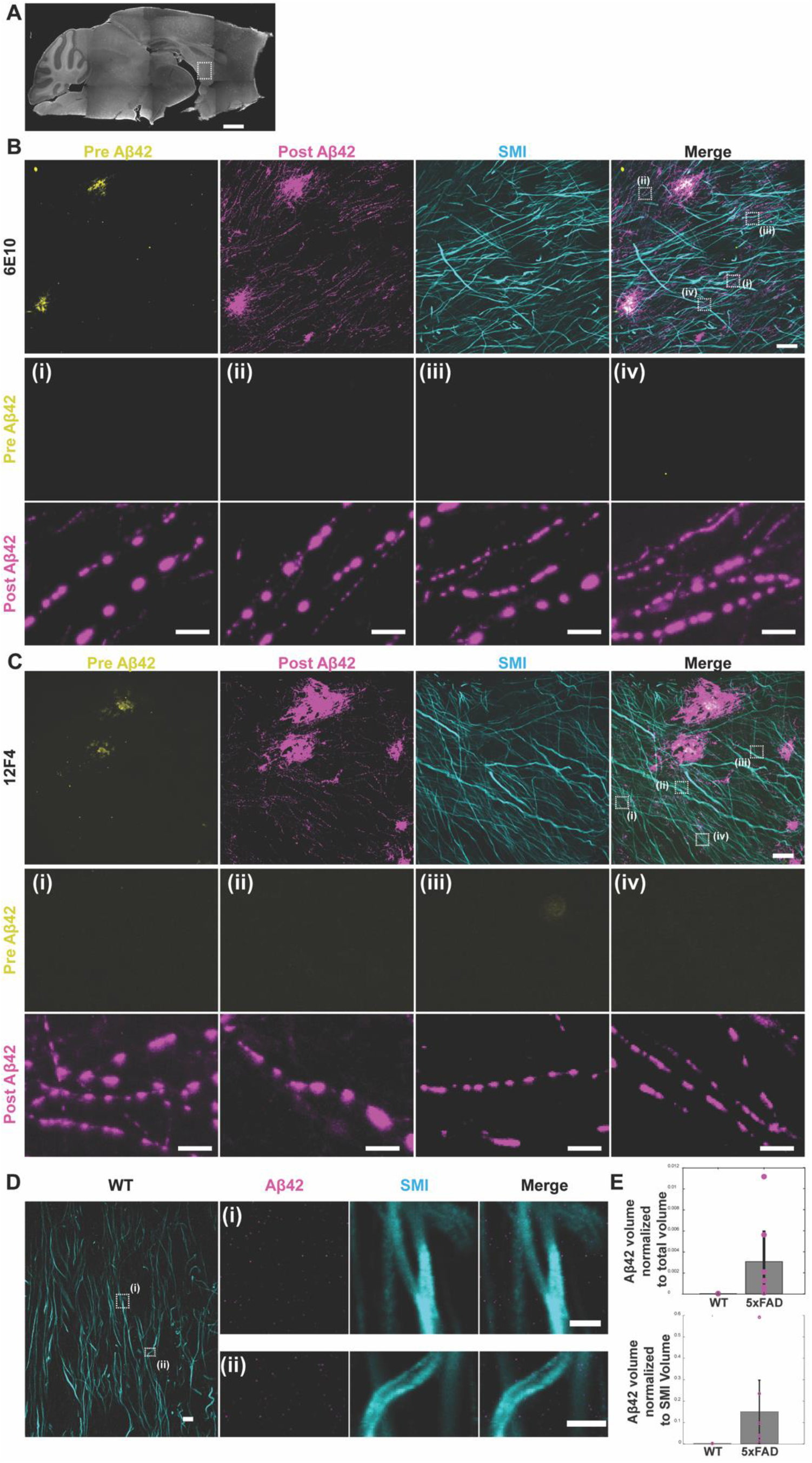
ExR reveals periodic nanoclusters of Aβ42 peptide in the fornix of Alzheimer’s model 5xFAD mice. (A) Epifluorescence image showing a sagittal section of a 5xFAD mouse brain with the fornix highlighted. (Scale bar, 1000 μm) (B-C) ExR confocal Images (max intensity projections) showing immunolabeling against Aβ42 peptide with two different monoclonal antibodies 6E10 (B) and 12F4 (C). From left to right: pre-expansion immunolabeling of Aβ42 (yellow), post-expansion labeling (magenta) of Aβ42, post-expansion SMI (neurofilament protein), and merged pre- and post-expansion staining of Aβ42 with post-expansion staining of SMI. Insets (i-iv) show regions of interest highlighted in the merged images in the fourth column; top panels, pre-expansion labeling; bottom panels, post-expansion labeling. Post-ExR staining reveals periodic nanostructures of β-amyloid, whereas pre-expansion staining can detect only large plaque centers. (Scale bar = 10 μm (upper panel images); 1 μm (bottom panels, i-iv)) (D) ExR confocal images showing post-expansion Aβ42 (magenta) and SMI (cyan) staining in wild-type (WT) mice. Left image, low magnification image in the fornix. Insets (i) and (ii), close-up views of Aβ42 and SMI staining patterns from boxed regions in the left image. (Scale bar = 4 μm (left panel); 1 μm (right panels, i-ii)) (E) Histograms showing the volume of Aβ42 clusters and aggregates in WT and 5xFAD specimens normalized to the total field of view (FOV) volume (top) or the volume of SMI (bottom). Two sample Kolmogorov-Smirnov test, p-values < 0.001. N= 14 data points (7 per condition) from 14 FOVs from 4 mice (2 mice per condition).

### Coclustering of amyloid nanodomains and ion channels

To understand the biological context of these periodic Aβ nanostructures, we stained brain slices with antibodies against the ion channels Nav1.6 and Kv7.2. In myelinated axons, Kv7 potassium channels and voltage gated sodium (Nav) channels colocalize tightly with nodes of Ranvier periodically along axons, while Nav1.6 and Kv7.2 are diffusely distributed along unmyelinated axons (*26*). Additionally, Alzheimer’s is associated with altered neuronal excitability and alterations in ion channels (*27*). We stained ExR-processed 5xFAD fornix-containing brain slices with antibodies against potassium channels (Kv7.2) and against β-amyloid (12F4) (Fig. 5A, Fig. S8). The periodic β-amyloid nanostructures colocalized with periodic nanoclusters of potassium channels. In ExR-processed WT fornix, such β-amyloid clusters, and clusters of potassium channel staining, were not found (Fig. S9). Co-staining of β-amyloid (12F4) and sodium channels (Nav1.6) showed complementary results (Fig. S10). To probe these periodicities and relationships further, we obtained a cross-sectional profile of a stretch of axon, showing that Aβ42 (magenta) and Kv7.2 (yellow) were highly overlapping and with similar periodicities (Fig. 5B). A further histogram analysis showed the periodicity of both repeated protein structures to be 500-1000 nm (Fig. 5C), which we confirmed by Fourier analysis (Fig. 5D-E). This periodicity is notably higher in spatial frequency than the inter-node-of-Ranvier distances of brain white matter, which range from 20-100μm (*28–30*). Our analysis along individual segments of axons also showed that a high fraction of Aβ42 clusters contained Kv7.2 clusters (Fig. 5F), a result that we confirmed by measuring the distance between the centroids of overlapping Aβ42 and Kv7.2 nanoclusters, finding extremely tight colocalization, within a few tens of nanometers (Fig. 5G). Median sizes of nanoclusters were around 150nm for both Aβ42 and Kv7.2 (Fig. 5H); indeed, overlapping nanodomains of Aβ42 and Kv7.2 were highly correlated in size (Fig. 5I), suggesting a potential linkage between how they are formed and organized as multiprotein complexes.

**Fig. 5.**
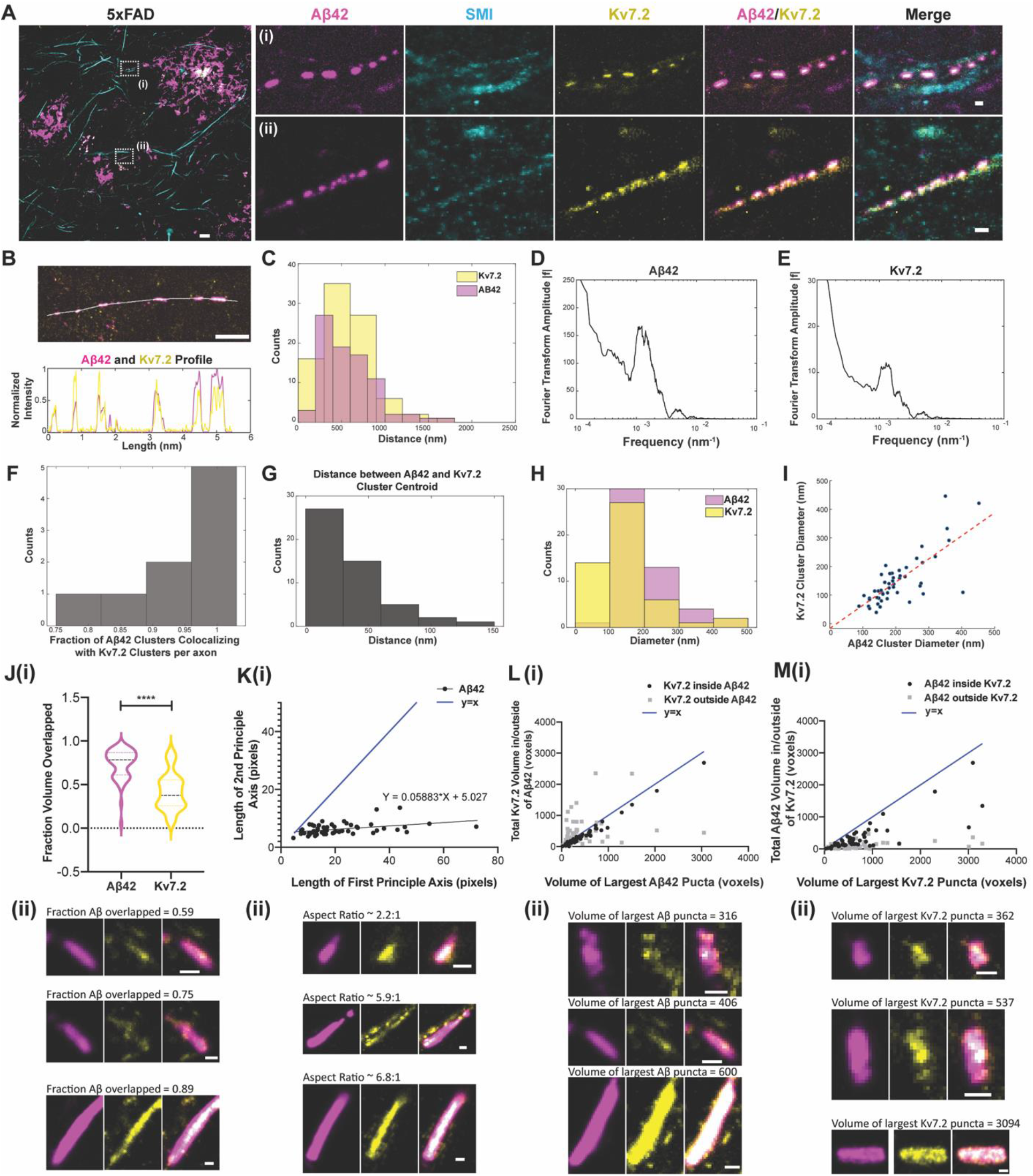
ExR reveals co-localized clusters of Aβ42 peptide and potassium ion channels in the fornix of Alzheimer’s model 5xFAD mice. (A) ExR confocal image (max intensity projections) showing post-expansion Aβ42 (magenta), SMI (cyan) and Kv7.2 (yellow) staining in the fornix of a 5xFAD mouse. Leftmost panel, merged low magnification image. (Scale bar = 4 μm); insets (i-ii) show close-up views of the boxed regions in the leftmost image for Aβ42, SMI, Kv7.2, Aβ42-Kv7.2 merged and Aβ42-Kv7.2-SMI merged respectively (scale bar = 400 nm; n = 5 fields of view from 2 slices from 2 mice) (B) ExR confocal image (max intensity projections) showing Aβ42 (magenta) and Kv7.2 (yellow) clusters in a 5xFAD mouse (top) with the indicated cross-section profile shown (bottom). (Scale bar = 1 μm) (C) Histograms showing distances between adjacent Aβ42 (magenta) and Kv7.2 (yellow) clusters in 5xFAD mice along imaged segments of axons (n=97 Aβ42 clusters, 92 Kv7.2 clusters from 9 axonal segments from 2 mice). (D-E) Fourier transformed plots of Aβ42 (d) and Kv7.2 (e) showing the same peak position (from the same data set as (C)). (F) Histogram showing the fraction of Aβ42 clusters colocalizing with Kv7.2 clusters along individual segments of axons (n=50 cluster pairs from 9 axonal segments from 2 mice). (G) Histogram showing the distance between the centroids of colocalized Aβ42 and Kv7.2 clusters (same data set as (F)). (H) Histograms showing the diameters of Aβ42 clusters (magenta) and Kv7.2 clusters (yellow) (n=50 cluster pairs from 9 axonal segments from 2 mice). (I) Scatter plot showing the diameters of colocalized Aβ42 and Kv7.2 clusters. (J)(i) Fraction of total Aβ42 and Kv7.2 puncta volume overlapped with one another in cropped Aβ42 clusters (n = 55 clusters from 2 5xFAD mice; p < 0.0001, pairwise t-test). ****P < 0.0001. (J)(ii) Representative images illustrating the difference in the proportion of mutually overlapped volume between Aβ42 and Kv7.2 as a fraction of total Aβ42 or Kv7.2 volume (scale bar = 100nm). (K)(i) Length of the second principal axis of the ellipsoid that has the same normalized second central moments as the largest Aβ42 punctum in an ROI, vs. the length of the first principal axis of this ellipsoid (in pixels, 1 pixel = 11.44 nm in x- and y-directions (transverse) and 26.67nm in the z-direction (axial); black points represent individual manually cropped ROIs; slope of best-fit line from simple linear regression = 0.05883, p = 0.0020, F = 10.59, df = 53). Compare to the line y = x (blue). (K)(ii) Representative images illustrating the oblong shape of Aβ42 puncta, for three first principal axis lengths: shorter (top row), medium (middle row) and longer (bottom row). While the length of the first principal axis varies significantly between these examples, the length of the second principal axis remains similar between the three clusters. (L)(i) Total volume (in voxels, 1 voxel = 11.44×11.44×26.67 or 3,490 nm3) of Kv7.2 puncta overlapped/inside (black, R^2^ = 0.980, p < 0.0001) and outside (gray, R^2^ = 0.0582, p = 0.0979) Aβ42 puncta, as a function of the volume of the largest Aβ42 puncta within an ROI, compared to the line y = x (blue). (m)(ii) Representative images illustrating that the volume of Kv7.2 outside of Aβ42 is relatively constant as Aβ42 puncta size increases. As in (K)(ii), the three clusters shown are ordered by increasing size. (M) The converse of (L): total volume of Aβ42 puncta overlapped/inside (black, R^2^ = 0.6531, p < 0.0001) and outside (gray, R^2^ = 0.1876, p = 0.0010) of Kv7.2 puncta, as a function of the volume of the largest Kv7.2 puncta within an ROI, compared to the line y = x (blue). (M)(ii). Representative images illustrating that the volume of Aβ42 outside of Kv7.2 is smaller than the volume of Aβ42 co-localized with Kv7.2, and both values are positively correlated with the volume of the largest Kv7.2 puncta. Scale bar of (J-M ii) = 100 nm.

To facilitate visualization of the 3D shapes of these Aβ42 and Kv7.2 puncta, we show three orthogonal slices in the x-y, x-z, and y-z planes (x- and y-directions being transverse directions, and the z-direction being axial) that intersect the center of each puncta. Qualitatively, we observed that the majority of these puncta are oblong, with smooth, continuous Aβ42 ellipsoids and more punctate Kv7.2 puncta, with some, but not all, Kv7.2 puncta found within Aβ42 puncta. A larger volume of Aβ42 (as a fraction of total Aβ42 puncta volume) was found inside Kv7.2 puncta compared to the fraction of Kv7.2 volume inside Aβ42 puncta (Fig. 5J). On average, the mean volume of an Aβ42 puncta was slightly larger than that of a Kv7.2 puncta (Fig. S11b, paired t-test, p = 0.0001, t=4.116, df=54). Quantification of shape characteristics confirmed these observations and revealed more subtle patterns. Representative images illustrating the observed trends are shown below each plot (Fig. 5Jii-nii). Despite the larger volume of Aβ42 puncta, the fraction of Aβ42 volume mutually overlapped with (inside of) Kv7.2 puncta was larger than the fraction of Kv7.2 mutually overlapped with (inside of) Aβ42 puncta (Fig. 5J; p < 0.0001, t=10.94, df=54). When considering the ellipsoidal shapes of these puncta, the relationship between the second and first principal axis lengths was highly sublinear (Fig. 5K), and the average aspect ratio (ratio of first to second principal axis length) was ~3.5:1 (mean 3.464, standard deviation 1.911, n = 55 puncta), indicating a highly oblong shape (slope of best-fit line from simple linear regression = 0.05883, p = 0.0020, F = 10.59, df = 53). While the number of Kv7.2 present in each manually cropped ROI was not correlated with the volume of the largest Aβ42 puncta within the cropped ROI (Fig. S11c; simple linear regression, 95% CI for slope [-0.003940, 0.005947]), the mean Kv7.2 puncta volume was significantly correlated with the mean Aβ42 puncta volume (Fig. S11d; simple linear regression, 95% CI of slope [0.09878-0.2437], R^2^ = 0.2977, p < 0.0001, F = 22.47, df = 53). Thus, as the size of Aβ42 aggregates increases, Kv7.2 aggregates increase in size, but not number. We found that the volume of Kv7.2 puncta inside Aβ42 puncta is highly correlated with the volume of the Aβ42 puncta (Fig. 5L; simple linear regression, 95% CI of slope [0.8353-9037], R^2^ = 0.98, p < 0.0001, F = 2,593, df = 53), but the volume of Kv7.2 puncta outside of Aβ42 puncta is not correlated to Aβ42 puncta volume (Fig. 5L; simple linear regression, 95% CI of slope [-0.03919, 0.4503], R^2^ = 0.05082, p = 0.0929, F = 2.838, df = 53). Conversely, the total volume of Aβ42 puncta inside of Kv7.2 puncta was not as strongly correlated (Fig. 5M; simple linear regression, 95% CI of slope [0.4198, 0.6307], R^2^ = 0.6531, p < 0.0001, F = 99.29, df = 53) with the volume of the largest Kv7.2 puncta in the cropped ROI. The volume of Aβ42 puncta outside of Kv7.2 was weakly but significantly correlated to the size of the largest Kv7.2 puncta (Fig. 5M; 95% CI of slope [0.0250, 0.09245] R^2^ = 0.1876, p = 0.0010, F = 12.24, df = 53). Finally, we found that the non-overlapped volume as a function of overlapped volume was much larger, on average, for Kv7.2 than for Aβ42 (Fig. S11e; paired t-test, p < 0.0001, t=5.985, df=54). Taken together, these results show that Kv7.2 and Aβ42 puncta are correlated in size only when physically colocalized in a tightly registered fashion, perhaps pointing to new potential hypothesized mechanisms of aggregation. While the biological relevance of this periodicity and colocalization of β-amyloid with Kv7.2 and Nav1.6 needs further investigation, it is interesting to note that, Nav1.6 and Kv7.2 ion channels can regulate neural excitability (*31–33*), and Aβ peptides have also been known to influence excitability (*34*). As these Aβ structures were often, but not always, colocalized with SMI-positive axons, and are highly reactive for Aβ42, we interpret these structures as periodic amyloid depositions. In the future, it will be interesting to see whether these structures play direct roles in neural hyperexcitability in Alzheimer’s.

It is curious to reflect here that periodicity and order (*35–45*) are often thought of as associated with healthy biological systems, whose functionality is supported by such crystallinity. On the other hand, disorder, misalignments and misfoldings are often tied to pathological states. Here we find a curious mixture of the two – a periodicity that seems to be associated with a pathological state, and that may have implications for new hypotheses related to Alzheimer’s pathology. We are excited to see how ExR might reveal many new kinds of previously invisible nanopatterns in healthy and disease states, due to its ease of use and applicability to multiple contexts, as here seen.

## Acknowledgments

We thank T. Biederer, C. Zhang and Y. Liu for antibody advice, S. Alon and K. Piatkevich for trainings and discussions and B. Kang for decrowding analysis advice. D. S. acknowledges funding from NIH K99/R00 Pathway to independence Award, M.E.S was supported by the National Science Foundation Graduate Research Fellowship under Grant No. 1745302, A.H.T. acknowledges funding from NARSAD. L.H.T. acknowledges funding from Ludwig Family Foundation, the JPB Foundation, NIH Grants RO1 NS102730 and RF1 AG054321, T.A.B acknowledges funding from NIMH R37MH080046, E.S.B. acknowledges funding from the Ludwig Family Foundation, the Open Philanthropy Project, John Doerr, Lisa Yang and the Tan-Yang Center for Autism Research at MIT, U. S. Army Research Laboratory and the U. S. Army Research Office under contract/grant number W911NF1510548, NIH Director’s Pioneer Award 1DP1NS087724, NIH R01MH110932, NIH R01EB024261, NIH U24NS109113, NIH R37MH08004613, NIH 1RM1HG008525, and the HHMI-Simons Faculty Scholars Program. Thanks to Brian Chow and Eric Betzig, as well as to many current and past members of the Synthetic Neurobiology group, for discussions.

The funders had no role in study design, data collection and analysis, decision to publish, or preparation of the manuscript.

## Supplementary Materials

Materials and Methods

Figures S1-S11

Tables S1-S8

Movies S1-S4

## Notes

### Competing Interest Statement

D.S., A.T.W., J.K., and E.S.B. are co-inventors on a patent application for ExR. E.S.B. is cofounder of a company seeking to deploy medical applications of ExM-related technologies.

