## Supplementary (Extended Data) Figures and Tables for "Expansion Revealing: Decrowding Proteins to Unmask Invisible Brain Nanostructures"

**This PDF file includes:**

Methods and Material

Fig. S1-11

Table. S1-7

**Content of Supplemental Information**

**Supplementary Figure 1.** Measurement of distortion of ExR in single z-slice confocal images of intact brain tissue vs. ExR tissue, imaging the same field of view before and after expansion, show that the expansion process is globally isotropic

**Supplementary Figure 2.** Comparison of antigen retrieval vs. ExR.

**Supplementary Figure 3.** Illustration and further quantification of synaptic decrowding analysis.

**Supplementary Figure 4.** Confocal images and comparison of pre-expansion and post-expansion immunostaining with antibodies against PSD95 (CST3450S).

**Supplementary Figure 5.** Analysis of distortion caused by post-expansion staining, as compared to classical pre-expansion staining.

**Supplementary Figure 6.** ExR confocal image showing immunolabeling of Aβ42 with two different monoclonal antibodies (a) D54D2 + 6E10 and (b) D54D2 + 12F4 with SMI staining in the fornix of 5xFAD mouse.

**Supplementary Figure 7.** Unexpanded tissue confocal image, a single z-slice, showing pre-expansion Aβ42 (yellow) and SMI (cyan) staining in the fornix of (a) WT and (b) 5xFAD mouse.

**Supplementary Figure 8.** Additional exemplary images of ExR confocal showing post expansion Aβ42 (magenta), SMI (cyan) and Kv7.2 (yellow) staining in the fornix of 5xFAD mouse.

**Supplementary Figure 9.** ExR confocal image showing post expansion Aβ42 (magenta), SMI (cyan) and Kv7.2 (yellow) staining in the fornix of WT mouse.

**Supplementary Figure 10.** ExR confocal image showing post expansion Aβ42 (magenta) and Nav1.6 (yellow) staining in the fornix of 5xFAD mouse.

**Supplementary Figure 11.** Additional exemplary images and analyses of Aβ42 and Kv7.2 nanocluster shapes and their relationship.

**Supplementary Table 1.** Numbers of technical replicates (synapses) used in Figure 2 analysis for each synaptic protein.

**Supplementary Table 2.** Statistical analysis for Figure 2i: synapse decrowding analysis statistics of mean signal amplitude within dilated reference channel ROIs.

**Supplementary Table 3.** Statistical analysis for Figure 2j: synapse decrowding analysis of mean puncta volume within dilated reference channel ROIs.

**Supplementary Table 4**. Accompanying statistics for Figure 2m-o and Supplementary Figure 5: analysis of distortion caused by post-expansion staining.

**Supplementary Table 5.** Statistical analysis for Figure 2k: synaptic puncta analysis for mean volume of the largest synaptic connected components.

**Supplementary Table 6.** Statistical analysis for Figure 2l: synaptic puncta analysis of mean signal-to-noise ratio within the largest synaptic connected components.

**Supplementary Table 7.** Statistical analysis for Supplementary Figure 2: comparison of the effects of antigen retrieval and ExR procedures on mean signal intensity within and outside of synaptic puncta.

**Supplementary Table 8.** Statistical analysis for Supplementary Figure 2: comparison of the effects of antigen retrieval and ExR procedures on total signal volume within and outside of synaptic puncta.

### Methods and Materials

#### Brain tissue preparation

All procedures involving animals were in accordance with the US National Institutes of Health Guide for the Care and Use of Laboratory Animals and approved by the Massachusetts Institute of Technology Committee on Animal Care. Both male and female wild type mice (C57BL/6 or Thy1-YFP) and 5xFAD mice were used, because of the current study's focus on developing and validating a novel technology. Mice were deeply anesthetized using isoflurane in room air. Mice were transcardially perfused at room temperature with ice cold 10 mL of 2 % (w/v) acrylamide in phosphate buffered saline (PBS) followed by ice cold 10 mL of 30 % acrylamide (w/v) and 4% paraformaldehyde in PBS. Brains were harvested and incubated in 20 mL of the same fixative solution (30 % acrylamide and 4% paraformaldehyde in PBS) at 4°C overnight. Fixed brains were transferred to 100 mM glycine at 4 °C for 6 h, then stored in PBS at 4°C for long term storage or sectioned to 50-100 μm-thick slices with a vibrating microtome (Leica VT1000S).

**Expansion of brain tissue slices**

For the 1st gelling step, brain slices were incubated in the 1st gelling solution (8.625% (w/v) sodium acrylate (SA), 2.5% (w/v) acrylamide (AA), 0.075% (w/v) N,N'-methylenebisacrylamide (Bis), 0.2% (w/v) ammonium persulfate (APS) initiator, 0.2% (w/v) tetramethylethylenediamine (TEMED) accelerator, 0.2% (w/v), 0.01% 4-Hydroxy-TEMPO (HT)) for 30 min at 4°C. Slices were embedded in 4 well dish gelling chambers on cover glasses surrounded by excess 1st gelling solution, and incubated at 37°C for 2 h. After the incubation, gels containing the tissue were cut out from the chamber and incubated with denaturation buffer (200 mM SDS, 200 mM NaCl, and 50 mM Tris pH 9) (*5*) for 1 h at 95°C. Denatured gels were fully expanded via 4 washes for 15 min each with 5 mL distilled (DI) water in a 6 well-plate.

For the re-embedding step, expanded 1st gels were incubated in re-embedding solution (13.75% (w/v) AA, 0.038% (w/v) Bis, 0.025% (w/v) APS, 0.025% (w/v) TEMED) twice, replacing the first solution with freshly made re-embedding solution for 1 h each time on a shaker at room temperature. The re-embedded gels were transferred to a 4 well dish gelling chamber on cover glasses and incubated with excess re-embedding solution for 2 h at 45°C. The re-embedded gels were washed 3 times for 15 min each with 5 mL PBS in a 6 well-plate.

For the 3rd gelling step, the re-embedded gels were incubated in 3rd gelling solution (8.625% (w/v) SA, 2.5% (w/v) AA, 0.038% (w/v) Bis, 0.025% (APS), 0.025% (w/v) TEMED) twice, replacing the first solution with fresh made 3rd gelling solution for 1 h each time on a shaker at room temperature. The 3rd gels were placed in a 4 well dish gelling chamber on cover glasses and incubated at 60 °C for 1 h.

Gels were fully expanded in DI water by changing excess water 5 times for 2 h each and trimming axially to reduce thickness to 1 mm to facilitate subsequent immunostaining and imaging.

**Immunostaining of expanded tissues**

Expanded gels were incubated in blocking solution (0.5% Triton X-100, 5% normal donkey serum (NDS) in PBS) for 2 h at room temperature. Gels were then incubated with primary antibodies (see Methods Antibody list) in '0.25T' blocking buffer (0.25% Triton X-100, 5% NDS in PBS) overnight at 4°C. Gels were washed in washing buffer (0.1% Triton X-100 in PBS) 6 times for 1 h each time on a shaker at room temperature. Gels were then incubated with secondary antibodies in blocking solution overnight at 4°C, and washed in washing buffer 6 times for 1 h each on a shaker at room temperature.

**Decrowding experiments**

<u>Fig 2B-H: Nanoscale-resolution imaging of synapses in somatosensory cortex</u>. For pre-expansion antibody staining, brain slices were incubated with primary antibodies in blocking solution overnight at 4°C. Stained tissues were washed in washing buffer 6 times for 1 hr each time on a shaker at room temperature. Secondary antibodies (100 μL, 1 mg/mL) were incubated with 6-((acryloyl)amino)hexanoic acid, succinimidyl ester (AcX) (2 μL, 1 mg/mL) overnight at room temperature to prepare AcX conjugated secondary antibodies. Primary antibody stained tissues were then incubated with AcX-secondary antibodies in blocking solution overnight at 4°C, and washed in washing buffer 6 times for 1 h each time on a shaker at room temperature. Tissue expansion was carried out the same as previously described. The tertiary antibodies were stained after expansion to bind against AcX-secondary antibodies to visualize the pre-expansion staining.

For post-expansion antibody staining, expanded gels were incubated with the same primary and secondary antibodies without AcX conjugation. Antibodies against Shank3 or Homer1 were provided as a reference channel after expansion.

**Quantification and validation of the decrowding effect**

<u>Fig. 2I-J. Decrowding analysis of manually segmented synapses.</u> We compared the amplitude of signal intensity in the foreground (putative synapses) and background (everything else). First, we manually identified, based on brightness and size of reference channel staining, 47-70 of the largest, brightest synapses per ~350x350x20 micron (physical units) field of view (**Table S1** for

exact numbers of synapses; one field of view per cortical layer, three cortical layers per sample, two mice per synaptic protein). We developed an automated method to segment putative synaptic puncta from background. First, background was subtracted from image stacks using ImageJ/Fiji's Rolling Ball algorithm with a radius of 50 pixels. Images were then binarized using a threshold calculated as seven times the standard deviation of the average intensity of manually-identified background regions selected every 10$^{th}$ slice of the z-stack. Binary images were passed through a 3-D median filter of radius 5x5x3 pixels to remove small puncta of non-specific staining. We then identified 3-D connected components from the filtered binary stack using MATLAB's "bwconncomp" function, with a pixel connectivity of 26, meaning that pixels are connected if their faces, edges, or corners touch. Connected components smaller than 120x120x120nm$^3$ (biological units) were removed, as most synapses are larger than this volume. ROIs, or putative synaptic puncta, were defined as 3-D connected components of the filtered, binary reference stack dilated using a disk structuring element with a radius of six pixels. A radius of six pixels (~100nm, in biological units) was chosen because both pre- and post-synaptic proteins of the same synapse, but not other synapses, fall within this range (the synaptic cleft is ~12-20nm (*46*), which we confirmed by manual inspection. Segmented synapses with zero filtered connected components (synaptic puncta) in the reference channel were excluded from further analysis. We calculated the average intensity of background-subtracted images either within or outside of these dilated reference ROIs (**Fig. 2I**) to measure signal increase within putative synapses relative to signal increase in the background. All images were acquired under the same microscope conditions to allow for comparison of mean signal intensity. The total volume of pre- or post-expansion staining test puncta located within dilated reference channel ROIs (**Fig. 2J**) was calculated from binarized stacks of pre- and post-expansion channels (thresholded and filtered as described for the reference channel) after multiplying the dilated binary reference stack (for inside dilated reference ROIs) or its inverse (for just outside dilated reference ROIs, but still within the manually-cropped synaptic area) by the binary pre- or post-expansion stack and calculating the sum of nonzero voxels for each product. Data are shown as the mean of each measure across the ~50 synapses per field of view, and the deviation was calculated as the standard error of these means across the six fields of view for each protein.

Fig. 2K-L. Quantification of synaptic properties. To compare volume and signal-to-noise ratio (SNR) of pre- and post-expansion staining, we used the same dataset for Fig. 2I, J analysis. First, the background was subtracted from image stacks using ImageJ/Fiji's Rolling Ball algorithm with a radius of 50 pixels. Images were then binarized using a threshold calculated as seven times the standard deviation of the average intensity of manually-identified background regions selected every 10$^{th}$ slice of the z-stack. We then identified and selected the biggest 3-D connected components in pre- and post-staining test channels separately in each layer of somatosensory cortex (L1, L2/3 and L4, respectively), as these are the most likely to be synapses. We calculated the voxel and signal intensity in the largest 3-D connected components from 49-70 manually selected synapses (**Table S1** for exact numbers for each layer, protein, and mouse). The signal intensity was divided by the standard deviation of the background intensity to calculate SNR.

Fig. 2M-N and Fig. S5: Analysis of distortion introduced by ExR relative to pre-expansion staining.

To calculate the number of synaptic puncta, background-subtracted images were first thresholded as described previously (based on a multiple of the standard deviation of manually-identified background regions) and passed through a 3x3x5 voxel (1 voxel = 17.16x17.16x40nm$^3$) median filter. MATLAB's "bwconncomp" function was used to find connected components (putative synaptic puncta, connectivity of 26), and connected components with fewer than 30 voxels of volume were excluded from further analysis. For the plots shown in **Fig. S5g-l**, images were shifted by one voxel in each direction and padded using the intensity values of the pixels that were shifted out at that step. To calculate pixel-wise correlations and autocorrelations (**Fig. S5g-h**), images were first normalized to their minimum and maximum intensity values. From these, we calculated the pairwise linear correlation coefficient (MATLAB's "corr") between pixel intensity values in the pre- and post-expansion staining channels, or pre-(/post-) and pre(/post)-expansion staining channels for autocorrelation. To calculate the half-maximal shift distance (**Fig. 2N-M**), we fit a third-degree polynomial (MATLAB's "fit" with "poly3") to the correlation or autocorrelation as a function of shift distance, and used the best-fit curve to estimate the shift distance at which the correlation or autocorrelation reached 50% of its maximum value. For the plots shown in **Fig. S5i-j,** the correlation was calculated with a slight modification to account for differences in puncta volume. First, background-subtracted images were masked based on the corresponding binary image. Second, nonzero pixels were divided by the mean intensity value in the nonzero regions. Finally, the correlation was calculated as the pairwise linear correlation coefficient (MATLAB's "corr") between masked, mean-normalized intensity values in the pre- and post-expansion staining channels. Mutually overlapped volume was calculated as the sum of nonzero pixels in intersection of the binary pre- and post-expansion staining z-stacks, and normalized to the total puncta volume (sum of nonzero pixels in the binary z-stack) in the pre-expansion staining channel (**Fig. S5k-l**). Each of these calculations was repeated for each shift in the x-, y-, and z-directions. Synapses with zero puncta in the pre- or post-expansion staining channels were excluded from analysis.

#### Comparison between antigen retrieval and decrowding effect

Fig. S2. Confocal images after immunostaining with antibodies against $Ca_v2.1$, PSD95 and Homer1 with or without antigen retrieval treatment to compare signal quality for antigen retrieval vs. ExR treatment. To determine whether antigen retrieval by heat denaturation alone is the dominant factor underlying increased signal quality afforded by ExR, we treated one group of tissues with a standard antigen-retrieval step (placing tissues in 20 mM sodium citrate at pH 8 and incubating at 100°C for 30 sec and 60°C for 30 min) (*11*). Tissues with or without this antigen-retrieval step were processed by ExR. Then, we compared the amplitude of signal intensity in foreground (putative synapses) and background (everything else). First, we manually identified, based on brightness and size of reference channel staining (Homer1 for Cav2.1, Shank3 for PSD95 and Homer1), 30 of the largest and brightest synapses per ~350x350x20 micron (physical units) field of view (n = 30 synapses from 1 field of view from 1 mouse). We used the automated segmentation procedure and calculated mean signal intensity and volume as described above (see "Decrowding analysis of manually segmented synapses").

**Protein distance measurement and Synaptic nanocolumn results analysis**

Fig 1H, Fig 3E-G, Fig 3I-P. For analysis, potential synapses were manually identified and selected based on 1) the juxtaposition of presynaptic clusters and postsynaptic clusters, and 2) the colocalization of clusters on the same side of the synapse. As camera pixel size was 167 nm (physical units) and the step size of the z-stack was 250 nm (physical units), the voxel size was not equivalent in all dimensions. Because isometric voxels were necessary for subsequent analysis, each voxel was then subdivided into 12 smaller isometric voxels, each 83.3 nm (physical units) in all three dimensions. For comparisons of RIM1/2 and PSD95, one cluster was shifted in space to optimally overlap with the other cluster, as previously described (*13, 47*). The vector of this shift was determined by cross-correlation of the two clusters, and defined both the transsynaptic axis and the distance between the two clusters. For comparisons of RIM1/2 and $Ca_v2.1$, the shift distance was set as 0, and for comparisons of RIM1/2 and PSD95, putative synapses with a RIM1/2 to PSD95 peak-to-peak distance of less than 20 or greater than 180 nm (biological units) were rejected from further analysis, consistent with the dimensions of the active zone and PSD. Any synapses that extended beyond the z-range of the imaged stack were also excluded.

The autocorrelation ($g_a(r)$) and the protein enrichment analyses were adapted from previously described localization data-based analyses (*13, 47*). The 3D autocorrelation function ($g_a(r)$) tests the general homogeneity of density within a defined volume. Because the function was normalized by the correlation of a homogenized object with the same shape and volume, homogeneous fluorescence within a synaptic cluster will give a $g_a(r)$ = 1 at all radii, and local intensity peaks will result in a $g_a(r)$ > 1 over a radius of the size of region of high intensity. The cluster boundary was defined based on fluorescent intensity after convolution with a spherical kernel (r ~ 300 nm).

The relative molecular distribution within two different protein clusters was characterized using a cross-enrichment analysis. The enrichment analysis was performed by measuring the angularly averaged voxel intensity as a function of the distance from the point of peak intensity in the reference channel, and then normalizing this value by the angularly averaged intensity (as a function of the distance from the point of peak intensity in the reference channel) based on an object of the same shape and volume as the real object with voxels set to the average intensity of the real object. The enrichment index was calculated by taking the average of the enrichment values within a radius of 60 nm from the peak of the reference channel.

Synapse numbers (n) for the analysis from 2 mice:

Autocorrelations

Cav2.1 (Fig. 3e): n = 144 synapses (Layer 1), 101 synapses (Layer 23), 103 synapses (Layer 4)

PSD95 (Fig. 3f): n = 144 synapses (Layer 1), 101 synapses (Layer 23), 103 synapses (Layer 4)

RIM1/2 (Fig. 3g): n = 144 synapses (Layer 1), 101 synapses (Layer 23), 103 synapses (Layer 4)

Enrichment analysis

RIM1/2 enrichment to PSD95 peak (Fig. 3i-j): n = 153 synapses (Layer 1), 103 synapses (Layer 23), 108 synapses (Layer 4)

PSD95 enrichment to RIM1/2 peak (Fig. 3k-l): n = 152 synapses (Layer 1), 102 synapses (Layer 23), 108 synapses (Layer 4)

$Ca_v$2.1 enrichment to RIM1/2 peak (Fig. 3m-n): n = 150 synapses (Layer 1), 103 synapses (Layer 23), 107 synapses (Layer 4)

RIM1/2 enrichment to $Ca_v$2.1 peak (Fig. 3o-p): n = 153 synapses (Layer 1), 99 synapses (Layer 23), 108 synapses (Layer 4)

Enrichment Index values (mean +/- S.D.):

RIM1/2 to PSD-95 peak (Fig. 3j): 1.585 +/- 0.330 (Layer 1), 1.535 +/- 0.358 (Layer 23), 1.545 +/- 0.332 (Layer 4)

PSD-95 to RIM1/2 peak (Fig. 3l): 1.611 +/- 0.308 (Layer 1), 1.632 +/- 0.269 (Layer 23), 1.622 +/- 0.285 (Layer 4)

$Ca_v$2.1 to RIM1/2 peak (Fig. 3n): 1.510 +/- 0.364 (Layer 1), 1.359 +/- 0.330 (Layer 23), 1.452 +/- 0.314 (Layer 4)

RIM1/2 to $Ca_v$2.1 peak (Fig. 3p): 1.493 +/- 0.330 (Layer 1), 1.317 +/- 0.311 (Layer 23), 1.422 +/- 0.322 (Layer 4)

**Alzheimer's results analysis**

Fig. 4E. Comparison of Aβ42 volume in WT vs 5xFAD. 3D image stacks of Aβ42 and SMI312 staining were background subtracted via rolling-ball background subtraction with a 200px radius using ImageJ/Fiji. For each color channel, the standard deviation for the background was calculated using a 75x75px window. Subsequently, each color channel was binarized by applying a threshold of 28 times the STD of the background. This value was determined by evaluating the amount of thresholding required to remove putative non-specific staining spots. Finally, after binarization, the volume of Aβ42 and SMI312 for each field of view (FOV) was determined by adding up the segmented pixels of each color channel.

Fig. 5C. Distance Measurement between Clusters. To calculate the distance between adjacent clusters for either Aβ42 or $K_v$7.2, clusters that line along SMI312 neurofilaments were manually cropped out in 3D. Then, after applying rolling-ball background subtraction with a 100px radius, the centroid of each cluster was annotated manually using ImageJ/Fiji in 3D. Given that the spacing between clusters is much larger than the size of each cluster, we reasoned that manual labeling of the centroids incurs minimal error. Finally, the distance between adjacent clusters was calculated in 3D.

Fig. 5G-I. Calculation of Aβ42 and $K_v$7.2 Cluster Diameter. After applying rolling-ball background subtraction with a 100px radius to 3D FOVs of Aβ42 and $K_V$7.2 staining, overlapping $K_V$7.2 and Aβ42 clusters were manually cropped out. After calculating the standard deviation of the background of each channel, the cropped images were binarized by applying a threshold ten times the standard deviation of the background. The volume of each cluster was then identified via Connected-Component analysis using MATLAB's "bwconncomp" function. Finally, the centroid and principal axis length of each cluster were determined using the associated "regionprops" function, which models each connected component region as an ellipsoid. The centroid values were then used to calculate the distance between overlapped Aβ42 and $K_v$7.2 clusters.

Fig. 5J-M. Aβ42 and Kv7.2 Cluster Shape Analysis. ROIs containing single Aβ42 puncta that were part of a periodic chain-like structure were manually identified (n = 55 ROIs, 5 ROIs per field of view, from 11 fields of view from two mice) from background-subtracted images (ImageJ/Fiji's Rolling Ball algorithm, radius of 50 pixels). To visualize the 3D shape of Aβ42 and $K_v$7.2 puncta within these ROIs, we resliced the image stack along both transverse dimensions, at equal spacing to the axial dimension. We display the middle slice in each stack in the x-y plane in **Fig. 5J(ii)-n(ii)** and the middle slice in each stack in the x-y, y-z, and x-z planes in **Fig. S11a** (where x- and y-directions are transverse, and z-direction is axial). To quantify shape features, we used CellProfiler's (*48*) Watershed (*49*) segmentation module to segment puncta within manually extracted ROIs, using a footprint of 30 pixels for each channel. A custom MATLAB script was deployed to calculate the number of puncta in each channel, mean and maximum volume and surface area of these puncta, length of the three principal axes of the ellipsoid that have the same normalized second central moments as the region for the largest puncta, and the total volume of puncta overlap between Aβ42 and $K_v$7.2 as the number of non-zero pixels in the intersection of the binary image stacks. To quantify the statistical significance of the relationships between these measures, either two-tailed paired t-tests or simple linear regression were used as described in the text.

**Expansion factor and root mean square error measurement**

Fig. S1. A Thy1-YFP mouse was perfused as described above and 50 µm coronal sections were prepared using a vibratome. Before expansion, YFP fluorescence was imaged in six fields of view from the cortex of three cortical slices. Subsequently, these slices were processed with the ExR protocol as described above. Expanded slices were then labeled with a primary antibody against GFP (thermo Fisher A-11122) and a secondary antibody (see Methods and Materials List of antibodies). The same fields of view imaged pre-expansion were identified and confocal images were acquired of the antibody staining. Pre and post ExR images were acquired on an Andor spinning disk (CSU-X1 Yokogawa) confocal microscope with a 40 × 1.15 numerical aperture water objective.

To determine distortion arising from the process of ExR, pre and post expansion images were aligned and deformations in images were determined as described previously (*3*). Briefly, pre and post ExR images were background subtracted with a Rolling Ball background subtraction algorithim (ImageJ/Fiji) with a 200 px radius. Then, corresponding confocal planes from pre and post images were identified and registered using Fiji's Turboreg method allowing for scaling and rigid rotation. Then, a custom MATLAB script was used to implement a B-spline based non-rigid registration between pre and post expansion images, yielding vector fields for deformation within the images. These vector fields were then used to calculate root-mean-square length distortions across varying lengths.

List of chemicals

| Product Name | Vendor | Product Number |
|---|---|---|

| | | |
|---|---|---|
| Sodium acrylate | Santa Cruz | CAS7446-81-3 |
| Acrylamide | Sigma | A9099 |
| N,N′-Methylenebisacrylamide (BIS) | Sigma | M7279 |
| Ammonium persulfate (APS) | Sigma | A3678 |
| N,N,N′,N′-Tetramethylethylenediamine (TEMED) | Sigma | T7024 |
| 4-Hydroxy-TEMPO (HT) | Sigma | 176141 |
| 6-((acryloyl)amino)hexanoic Acid, Succinimidyl Ester (AcX) | Thermo Fisher | A20770 |
| Sodium dodecyl sulfate (SDS) | Sigma | 436143 |
| Sodium Chloride (NaCl) | Thermo Fisher | AM9760 |
| Tris Buffer | Fisher scientific | 77-86-1 |
| Paraformaldehyde | Electron Microscopy Sciences | 15710 |
| Triton X-100 | Sigma | X100 |
| Glycine | Sigma | 50046 |
| PBS 10x | Thermo Fisher | 70011044 |
| Normal Donkey Serum | Jackson ImmunoResearch | 017-000-121 |
| Sodium citrate dihydrate | Sigma | W302600 |

List of antibodies

| Primary / Secondary | Target | Host | Vendor | Product number | Dilution |
|---|---|---|---|---|---|
| Primary | Cav1.2 | Guinea pig | Synaptic Systems | 152 205 | 1:200 |
| Primary | RIM1/2 | Rabbit | Synaptic Systems | 140 203 | 1:200 |
| Primary | PSD95 | Mouse | Thermo Fisher | MA1-046 | 1:200 |
| Primary | PSD95 | Rabbit | Cell Signaling Technology | CST3450S | 1:200 |
| Primary | SynGAP | Rabbit | Thermo Fisher | PA1-046 | 1:200 |
| Primary | Homer1 | Rabbit | Synaptic Systems | 160 003 | 1:200 |
| Primary | Bassoon | Rabbit | Synaptic Systems | 141 003 | 1:200 |
| Primary | Shank3 | Guinea pig | Synaptic Systems | 162 304 | 1:200 |
| Primary | Aβ42 (6E10) | Mouse | BioLegend | SIG39320 | 1:200 |
| Primary | Aβ42 (12F4) | Mouse | BioLegend | SIG39142 | 1:200 |
| Primary | Aβ42 (D54D2) | Rabbit | Cell Signaling Technology | CST8243S | 1:200 |
| Primary | SMI | Chicken | Abcam | ab4680 | 1:400 |
| Primary | Kv7.2 | Mouse | Santa Cruz | sc-271852 | 1:200 |
| Primary | Nav1.6 | Rabbit | Abcam | ab65166 | 1:200 |
| Secondary | Mouse | Goat | ThermoFisher | A28175 (Alexa Fluor 488 nm) | 1:200 |

| | | | | | |
|---|---|---|---|---|---|
| Secondary | Mouse | Goat | ThermoFisher | A11031 (Alexa Fluor 546 nm) | 1:200 |
| Secondary | Mouse | Donkey | Biotium | 20124 (CF 633 nm) | 1:200 |
| Secondary | Mouse | Donkey | ThermoFisher | A10036 (Alexa Fluor 546 nm) | 1:200 |
| Secondary | Rabbit | Goat | ThermoFisher | A11034 (Alexa Fluor 488 nm) | 1:200 |
| Secondary | Rabbit | Goat | ThermoFisher | A11035 (Alexa Fluor 546 nm) | 1:200 |
| Secondary | Rabbit | Donkey | Biotium | 20125 (CF 633 nm) | 1:200 |
| Secondary | Rabbit | Donkey | ThermoFisher | A10040 (Alexa Fluor 546 nm) | 1:200 |
| Secondary | Guinea pig | Donkey | Biotium | 20171 (CF 633 nm) | 1:200 |
| Secondary | Chicken | Goat | ThermoFisher | A11039 (Alexa Fluor 488 nm) | 1:200 |
| Secondary | Chicken | Donkey | Biotium | 20168 (CF 633 nm) | 1:200 |

Monomer solution of ExR

| Component | Stock Concentration* | Amount (mL) | Final Concentration* |
|---|---|---|---|
| Sodium acrylate | 38 | 9 | 8.6 |
| Acrylamide | 50 | 2 | 2.5 |

| | | | |
|---|---|---|---|
| Sodium Chloride | 29.2 | 16 | 11.7 |
| PBS | 10x | 4 | 1x |
| Water | - | 3.6 | - |
| Total | - | 34.6 | - |

* All concentrations in g/100 mL except PBS

Gel solution of ExR

| Chemical | Stock Concentration (g/100 mL) | 1st gel solution (μL) | Re-embedding solution (μL) | 3rd gel solution (μL) |
|---|---|---|---|---|
| Monomer | - | 864 | - | 864 |
| Acrylamide | 50 | - | 275 | - |
| Bis acrylamide | 2 | 40 | 18.75 | 20 |
| Water | - | 36 | 701.25 | 111 |
| 4-HT | 0.5 | 20 | - | - |
| TEMED | 10 | 20 | 2.5 | 2.5 |
| APS | 10 | 20 | 2.5 | 2.5 |
| Total (mL) | - | 1 | 1 | 1 |

**ExR Procedure**

1. Mouse tissue slices

i. Anesthetize mice using isoflurane in oxygen and perfuse with 10 mL of 2% acrylamide in PBS followed by 10 mL of 30% acrylamide and 4% paraformaldehyde in PBS.

ii. Harvest brains and incubate in 20 mL of the same fixative solution (30% acrylamide and 4% formaldehyde in PBS) at 4°C overnight.

iii. Transfer fixed brains to 100 mM Glycine at 4°C for 6 h.

iv. Store tissues in PBS at 4°C for long term storage.

v. Slice tissues on a vibrating microtome to a thickness of 50-100 μm.

2. Gellation

A. Gelling for 1st expansion

i. Incubate brain slices in the 1st gelling solution for 30 min at 4°C.

ii. Place brain slices with excess 1st gelling solution between two #1.5 coverglass separated by two pieces of #1.5 coverglass, and then incubate at 37°C for 2 h.

iii. Cut out gels from the chamber and incubate with denaturation buffer (200 mM SDS, 200 mM NaCl, and 50 mM Tris pH 9) for 1 h at 95°C.

iv. Wash gels 4 times with DI water in shaker and expand gels in DI water at 4°C overnight.

B. Re-embedding

i. Incubate expanded 1st gels in re-embedding solution twice for 1 h each time in shaker at room temperature.

ii. Transfer gels between #1.5 coverglass separated by slide glass and incubate with excess re-embedding solution at 45°C for 2 h.

iii. Wash gels 3 times with PBS in shaker.

C. 3rd gelling

i. Incubate the re-embedded gels in the 3rd gelling solution twice for 1 h each time in shaker at room temperature.

ii. Transfer gels between #1.5 coverglass separated by slide glass and incubate at 60°C for 1 h.

iii. Wash gels 4 times with DI water in shaker and expand gels in DI water at 4°C overnight.

iv. Trim gels axially to 1 mm thickness.

3. Staining

i. Incubate gels in blocking solution (0.5% Triton X-100, 5% normal donkey serum (NDS) in PBS) for 2 h at room temperature.

ii. Incubate gels with primary antibodies in ‘0.25T’ blocking buffer (0.25% Triton X-100, 5% NDS in PBS) overnight at 4°C.

iii. Wash gels with washing buffer (0.1% Triton X-100 in PBS) 6 times for 1 h each time.

iv. Incubate gels with secondary antibodies in blocking solution at 4°C overnight.

v. Wash gels with washing buffer (0.1% Triton X-100 in PBS) 6 times for 1 h each time and expand gels in DI water for 20x expansion or 0.05x PBS for 15x expansion.

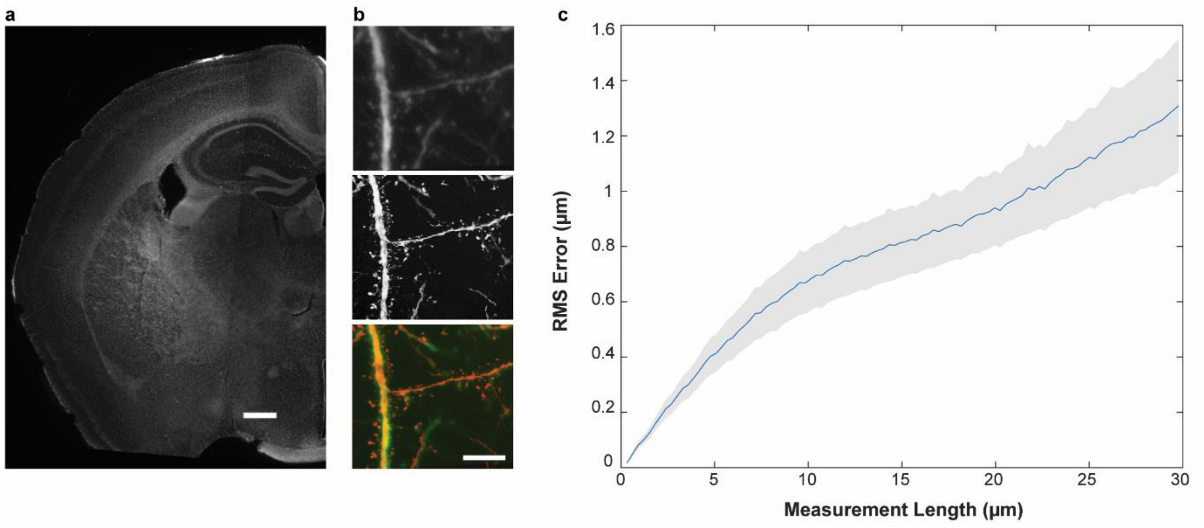


**Fig. S1.** Measurement of distortion of ExR in single z-slice confocal images of intact brain tissue vs. ExR tissue, imaging the same field of view before and after expansion, show that the expansion process is globally isotropic. (a) Widefield image of Thy1-YFP brain slice before expansion. (Scale bar, 500 µm.) (b) Confocal images after immunostaining with antibodies against GFP before expansion (top), after expansion (middle) and merged (bottom). (Scale bar, 5 µm, in biological units, meaning physical size divided by the expansion factor, here and throughout the paper unless otherwise indicated) (c) Root mean square error, vs. measurement length (in biological units), calculated via a non-rigid registration algorithm of intact vs. ExR-processed brain (n=6 fields of view from 3 coronal tissue sections, from one mouse). Blue line, mean; gray shading, standard deviation.

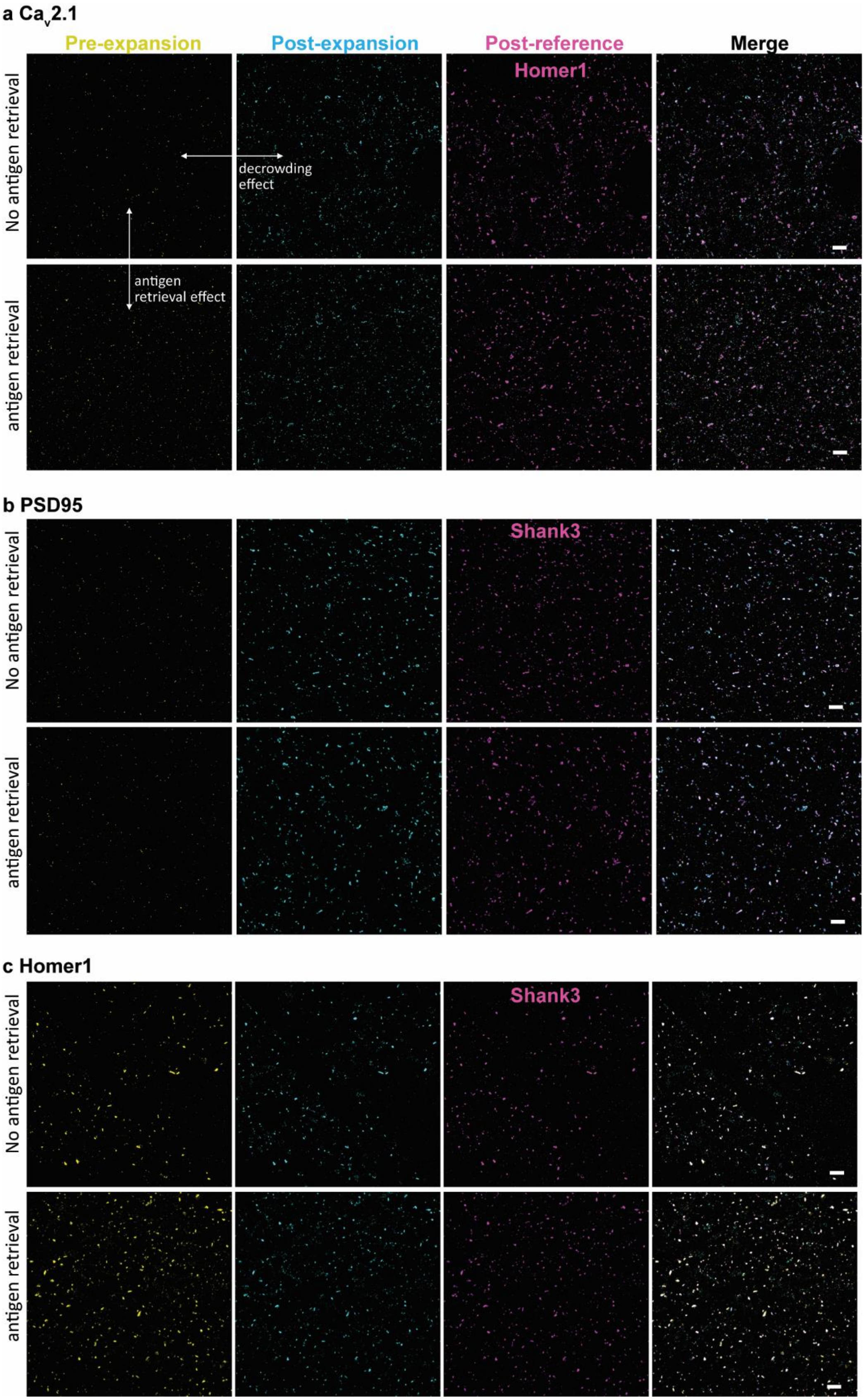
a Cav2.1
Pre-expansion
Post-expansion
Post-reference
Merge
Homer1
No antigen retrieval
antigen retrieval
decrowding effect
antigen retrieval effect
b PSD95
Shank3
No antigen retrieval
antigen retrieval
c Homer1
Shank3
No antigen retrieval
antigen retrieval

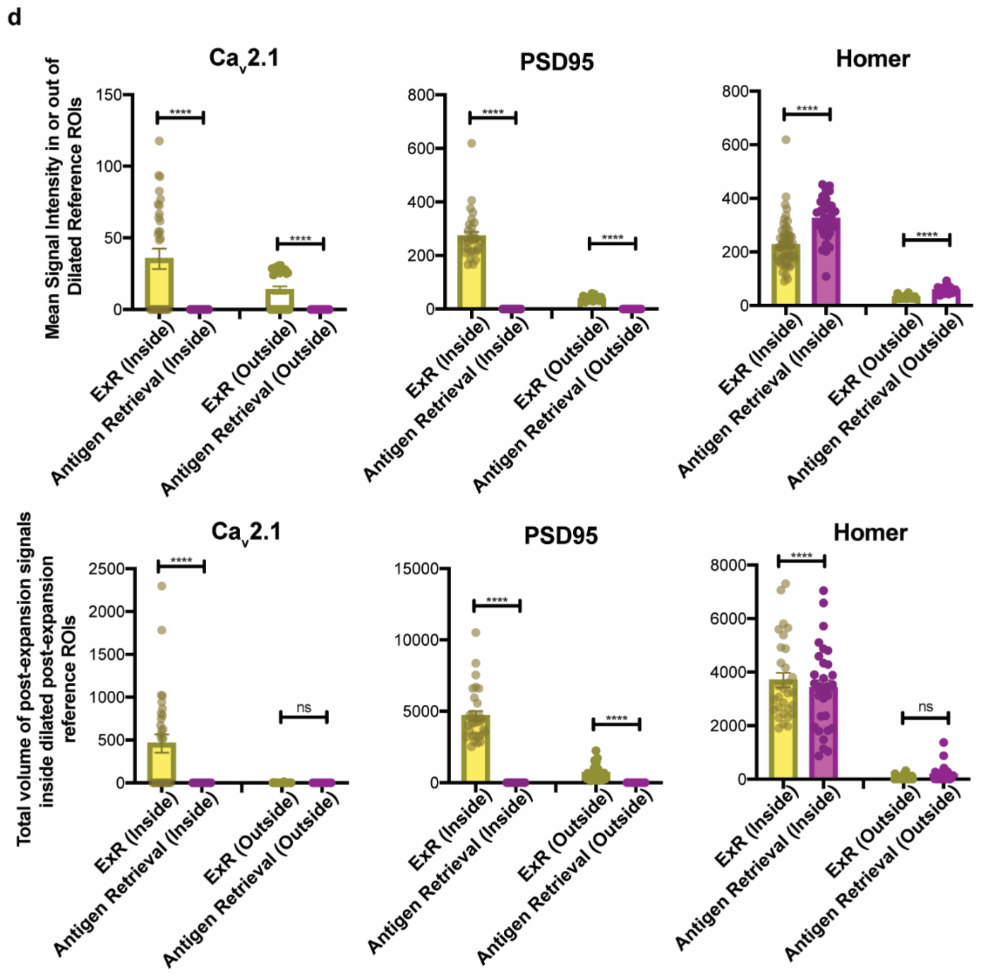


**Fig. S2.** Comparison of antigen retrieval vs. ExR. **(a-c)** Confocal images (max intensity projection) of pre-expansion and post-expansion immunostaining using the same imaging setup with antibodies against **(a)** Ca$_V$2.1, (**b**) PSD95, and **(c)** Homer1 with or without antigen retrieval. Scale bar, 1 μm. **(d)** Quantification of mean signal intensity (top row; all images were acquired under the same exposure and laser power) and total volume (bottom row) in or out of dilated reference ROIs with or without antigen retrieval steps (Sidak's multiple comparisons test following ANOVA; ****, $P < 0.0001$; ns, not significant). Antigen retrieval does not improve staining of proteins (Ca$_V$2.1 and PSD95) that cannot be stained without decrowding (ExR). In the case of Homer1, which shows good staining quality even without decrowding, antigen retrieval slightly increased overall mean signal intensity in both foreground and background, but did not increase volume. Thus, for some antibodies, the traditional antigen retrieval procedure may increase overall signal intensity relative to decrowding (ExR), potentially in a non-specific manner. We conclude that the decrowding effect due to expansion, not antigen retrieval, is the dominant factor contributing to increased staining quality with ExR. For full statistics, see **Tables S7** and S**8**.

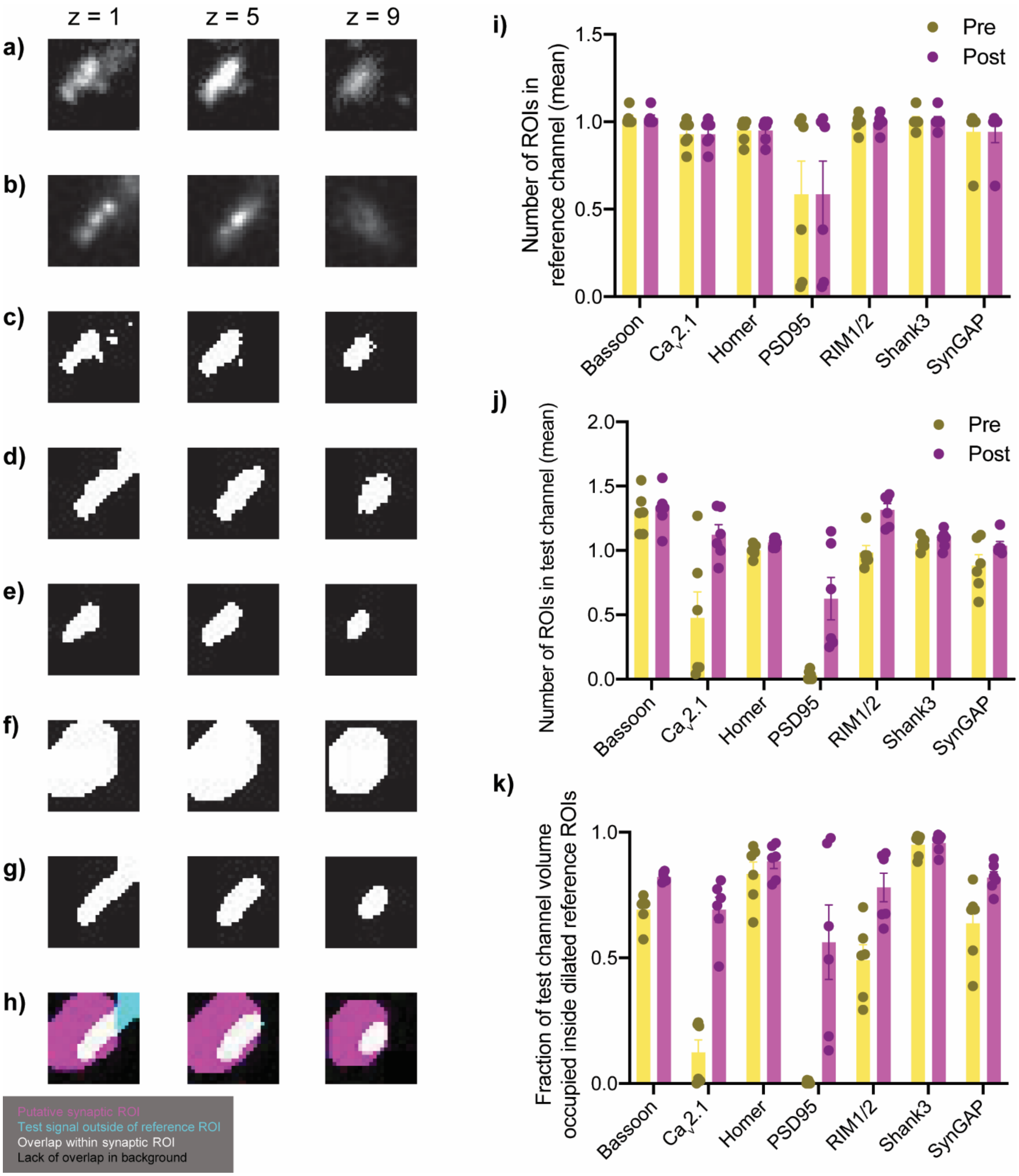


**Fig. S3.** Illustration and further quantification of synaptic decrowding analysis. Synapse size = 25x25x11 voxels ~ 425x425x440nm. Test (post-expansion) stain is Bassoon, reference stain (also post-expansion) is Homer1. **(a)** Background subtracted reference, **(b)** Background subtracted test stain, **(c)** Binary reference, **(d)** Binary test, **(e)** Filtered binary reference, **(f)**

Dilated reference, **(g)** Filtered binary test stain, and **(h)** Overlay of (**f**) and (**g**). **(i)** Mean number of ROIs per synapse for reference channel, for various test channels, shown on the x-axis. **(j)** Mean number of ROIs per synapse for pre- or post-expansion test channels. **(k)** Fraction of test channel (g) volume occupied inside dilated post-expansion reference ROIs (f). This measure can be interpreted as a true positive rate, because it is the fraction of "true" synapses that we are able to detect with ExR.

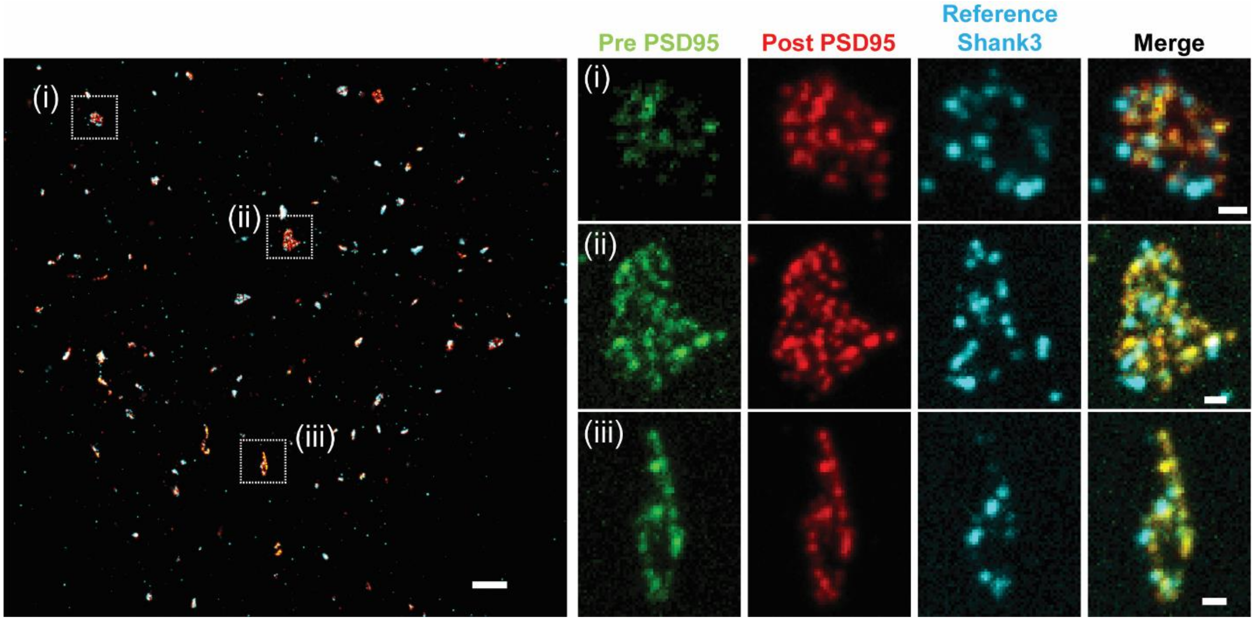


**Fig. S4.** Confocal images (single z-slice) of somatosensory cortex layer 2/3, comparing pre-expansion and post-expansion immunostaining with antibodies against PSD95 (from Cell Signaling Technology, product number CST3450S) and post-expansion Shank3 for the reference (n = 3 fields of view of 2 slices from 1 mouse). Scale bar, 1 μm (left panel), 100 nm (right panel).

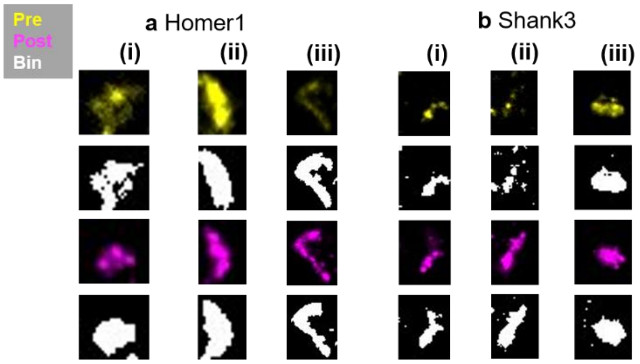


c

| | Homer1 | | | Shank3 | | |
|---|---|---|---|---|---|---|
| | a(i) | a(ii) | a(iii) | b(i) | b(ii) | b(iii) |
| # pre | 1 | 1 | 2 | 3 | 1 | 1 |
| # post | 1 | 1 | 1 | 2 | 1 | 1 |

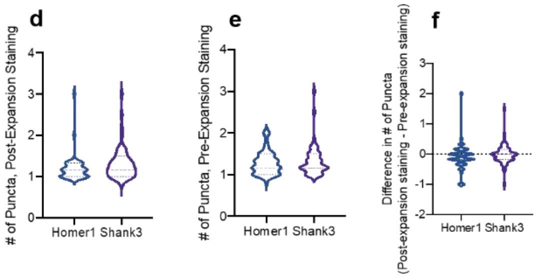


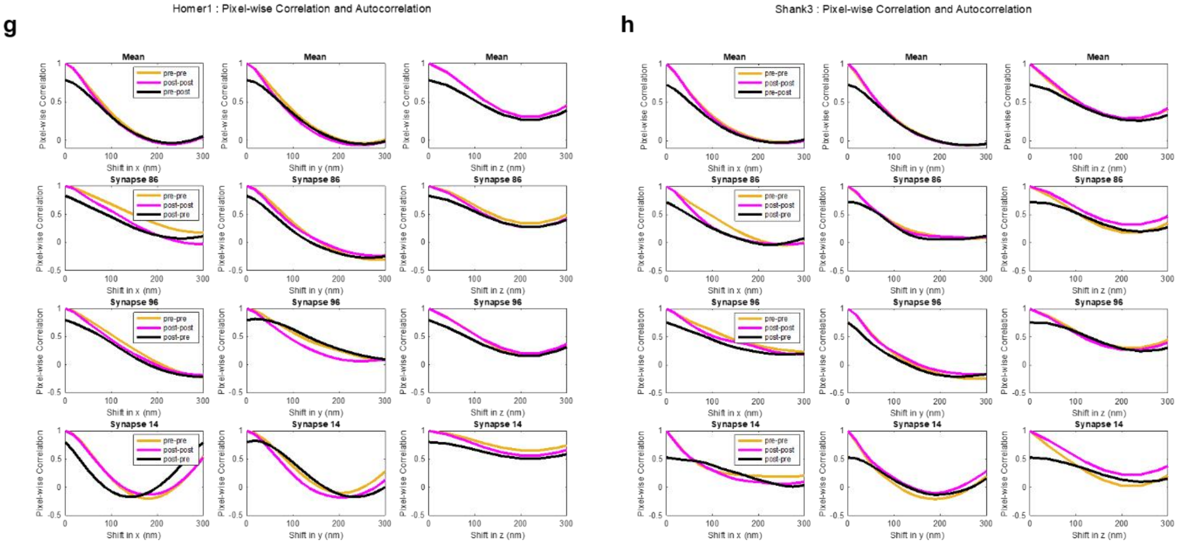

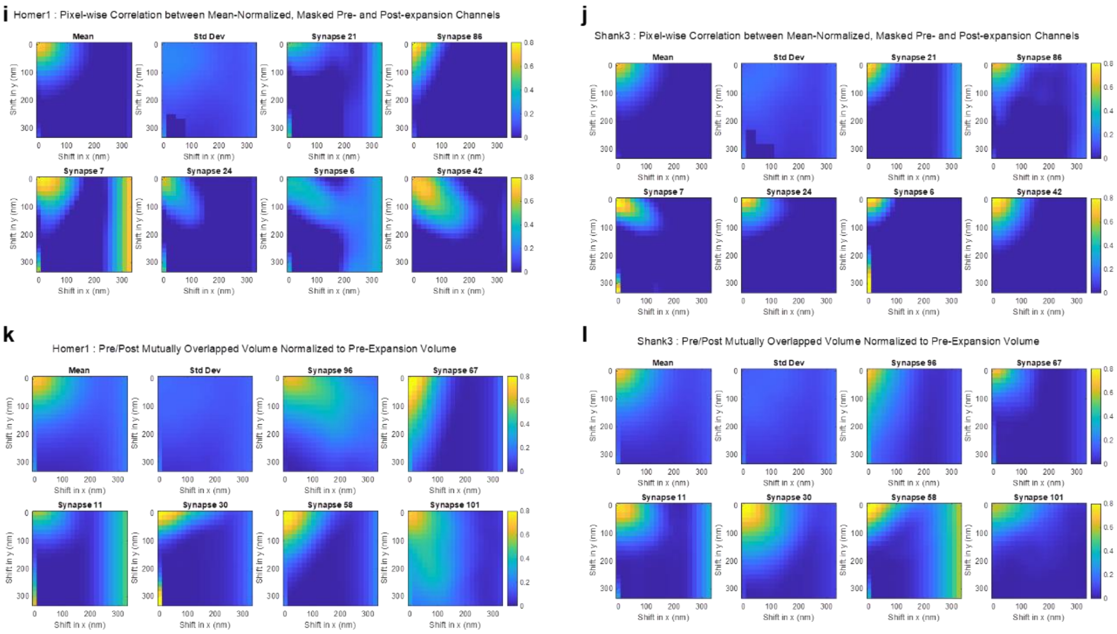


**Fig. S5.** Analysis of distortion caused by post-expansion staining, as compared to classical pre-expansion staining. **(a-b)** Representative background-subtracted and binary images of Homer1 **(a)** and Shank3 **(b)** in pre- and post-expansion staining channels (top row (yellow): pre-expansion channel, second row (black/white): binary pre-expansion channel, third row (magenta): post-expansion channel, bottom row (black/white): binary post-expansion channel). **(c)** Number of synaptic puncta for pre- and post-expansion staining channels, after filtration, for the images in **(a-b)**. **(d)** Population distribution (violin plot of density, with a dashed line at the median and dotted lines at the quartiles) of the number of synaptic puncta in the pre-expansion staining channel for Homer1 and Shank3 (see **Table S4** for statistics for this figure). **(e)** Population distribution of the number of synaptic puncta in the post-expansion staining channel for Homer1 and Shank3. **(f)** Difference in the number of synaptic puncta between post- and pre-expansion staining channels normalized to the number of synaptic puncta in the pre-expansion staining channel. **(g-h)** Pixel-wise autocorrelation between pre-expansion (pre-pre, yellow), post-expansion (post-post, magenta), and pixel-wise correlation between pre- and post-expansion (pre-post, black) as a function of shift distance in x- (left column), y- (middle column), and z- (right column) directions for Homer1 **(g)** and Shank3 **(h)**. The mean across all synapses is shown in the top row, and representative synapses are shown in the second through fourth rows. These values were used to calculate the linearized error measure shown in **Fig 2n-o**. **(i-j)** Pixel-wise correlation between mean-normalized, masked pre- and post-expansion channels as a function of shift distance in x- and y-directions (z=1) for Homer1 **(i)** and Shank3 **(j)**. The mean across all synapses is shown in the top left, standard deviation across all synapses shown in second from the top left, and representative synapses are shown in the remaining plots. **(k-l)** Mutually overlapped volume between pre- and post-expansion stained synaptic puncta, normalized to total puncta volume in the pre-expansion staining channel, as a function of shift distance in x- and y-directions (z=1) for Homer1 **(k)** and Shank3 **(l)**. The mean across all synapses is shown in the top left, standard deviation across all synapses shown in second from the top left, and representative synapses are shown in the remaining plots. Analysis was conducted on 330 (before exclusion based on size) synapses for Shank3, and 315 (before exclusion based on size) synapses for Homer1 from two mice (see **Table S4** for exact numbers).

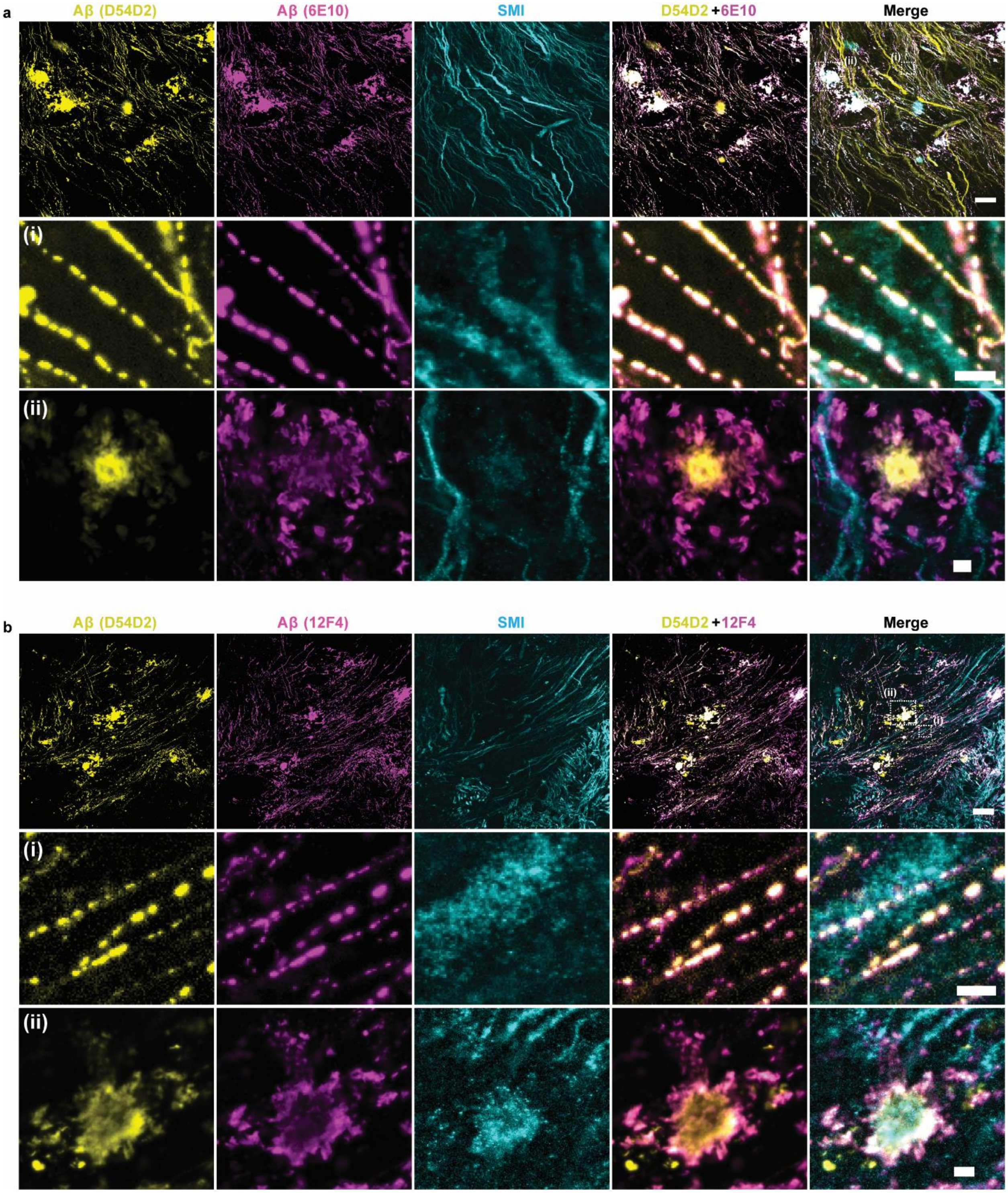


**Fig. S6.** ExR confocal images (single z-slices) showing immunolabeling of Aβ42 with two different monoclonal antibodies (a) D54D2 + 6E10 and (b) D54D2 + 12F4 with SMI co-staining in the fornix of 5xFAD mouse (n=3 fields of view of 2 slices from 2 mice). Scale bar, 10 μm (top row), 1 μm (i, ii panels).

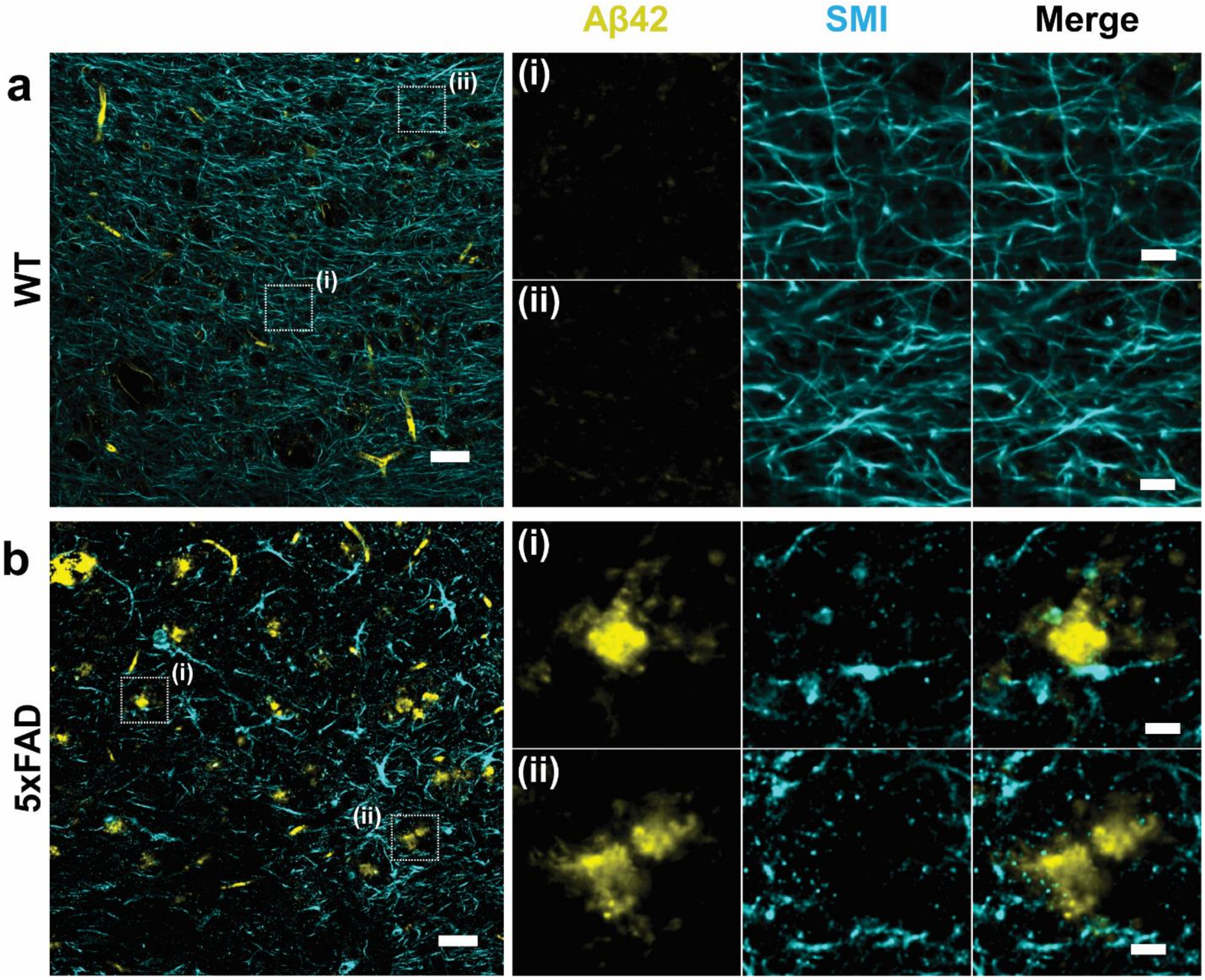


**Fig. S7.** Unexpanded tissue confocal image, a single z-slice, showing pre-expansion Aβ42 (yellow) and SMI (cyan) staining in the fornix of (a) WT and (b) 5xFAD mice (n=3 fields of view of 1 slice from 1 mouse per WT and 5xFAD, respectively). Scale bar, 30 μm (left panel) and 6 μm (panels i, ii)).

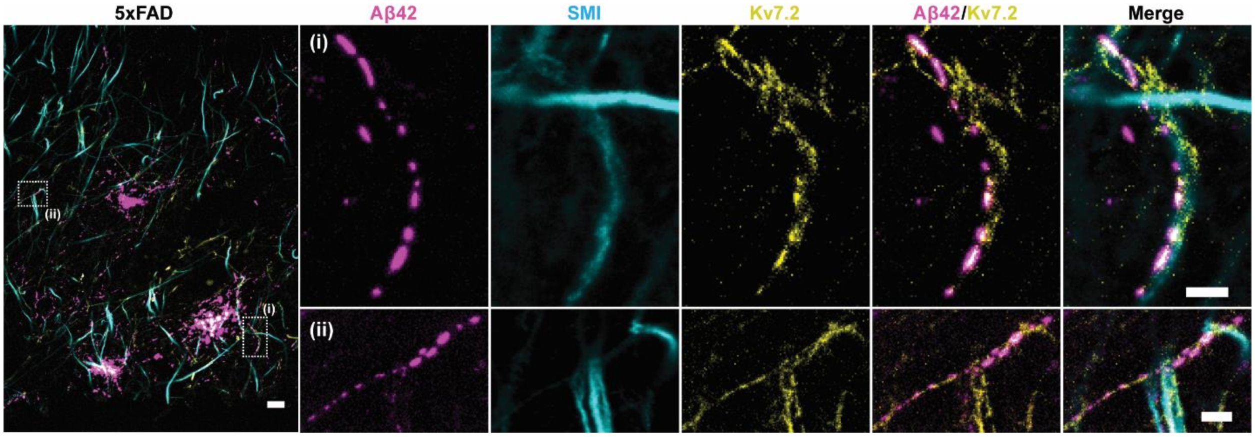


**Fig. S8.** Additional exemplar (i.e., with large puncta for easy visibility) confocal ExR images (single z-slice) showing post expansion Aβ42 (magenta), SMI (cyan) and Kv7.2 (yellow) staining in the fornix of 5xFAD mouse (n=3 fields of view of 2 slices from 2 mice). Scale bar = 4 μm (left panel), 400 nm (panels i, ii).

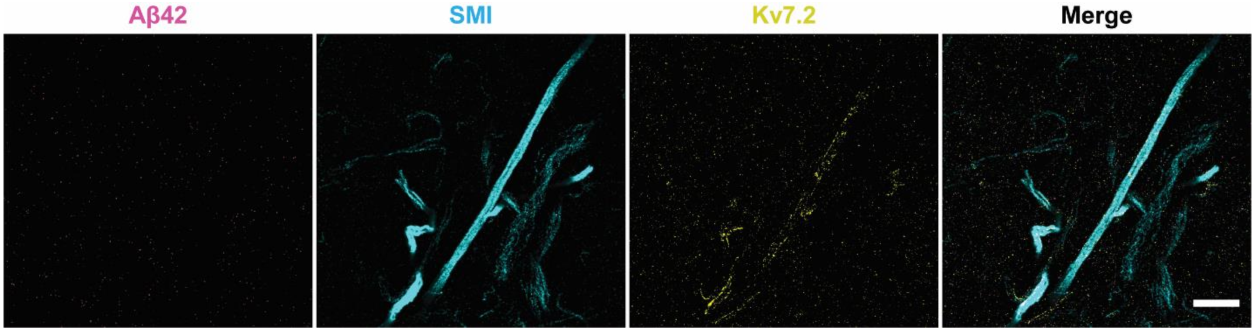


**Fig. S9.** ExR confocal image (single z-slice)  showing post expansion Aβ42 (magenta), SMI (cyan) and Kv7.2 (yellow) staining in the fornix of WT mouse (n=3 fields of view of 1 slice from 1 mouse). Scale bar = 2 μm.

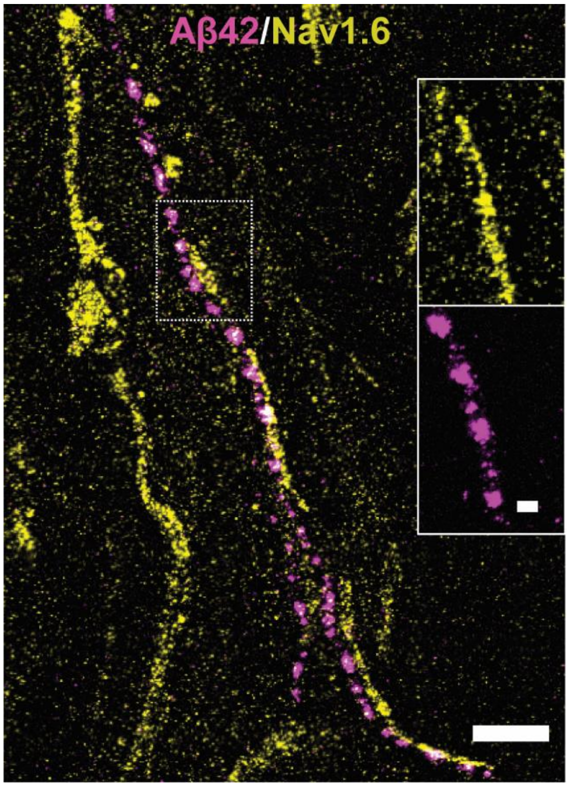


**Fig. S10.** ExR confocal image (single z-slice) showing post expansion Aβ42 (magenta) and Nav1.6 (yellow) staining in the fornix of 5xFAD mouse (n=3 fields of view of 1 slice from 1 mouse). Scale bar = 1 μm; 200 nm (inset).

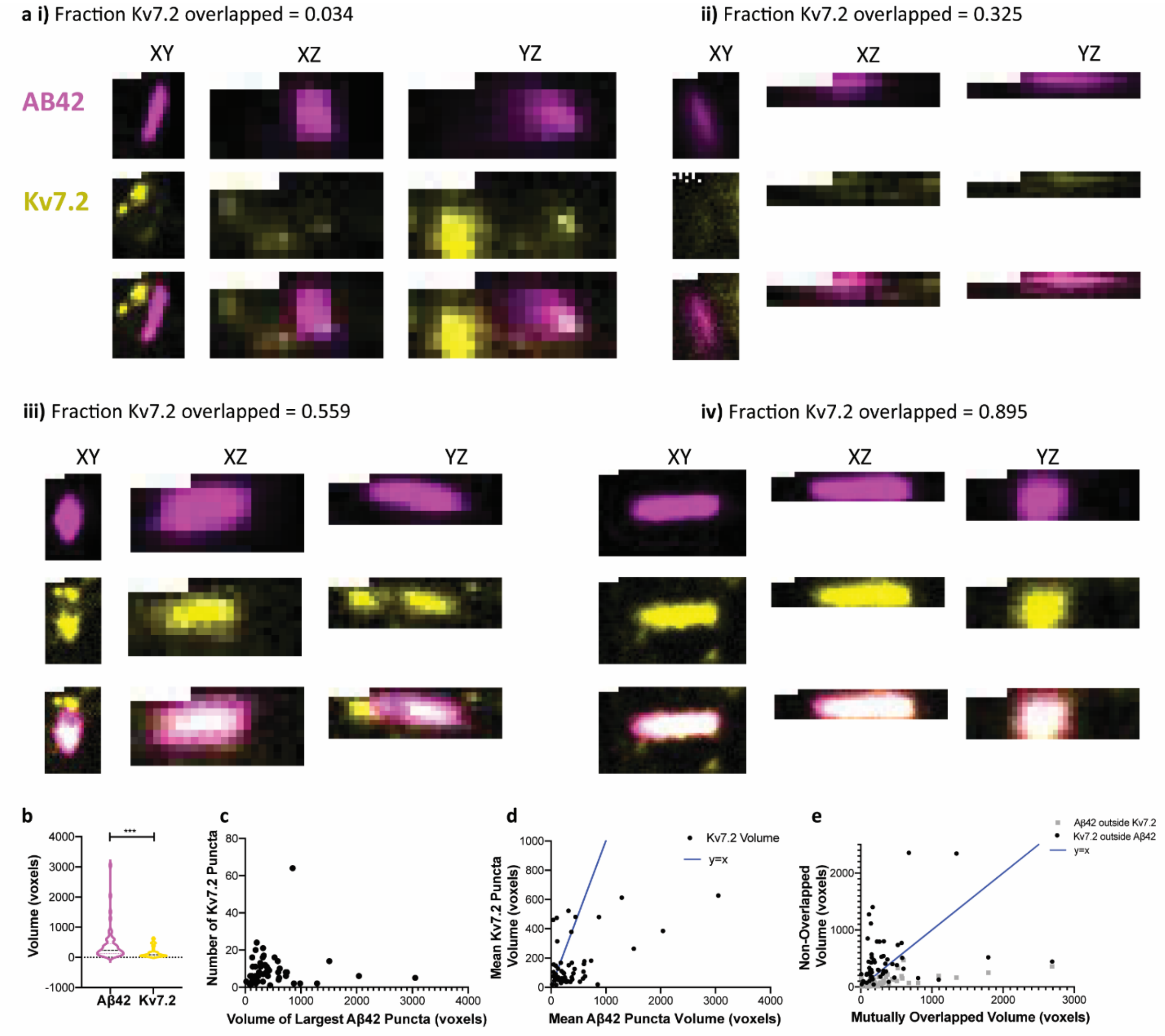


**Fig. S11.** Additional exemplary (good quality, not necessary representative) images and analyses of Aβ42 and $K_v$7.2 nanocluster shapes and their relationships. (**a**) 3D representation of an Aβ42 and $K_v$7.2 nanocluster using orthogonal slices in the x-y, x-z, and y-z planes, at the center of each nanocluster for nanoclusters with 3.4% **(i)**, 32.5% **(ii)**, 55.9% **(iii)**, and 89.5% **(iv)** of total $K_v$7.2 volume inside the Aβ42 puncta. The orthogonal projections illustrate the oblong shape of these nanoclusters. Scale bar = 100 nm. (**b**) Distribution of mean puncta volume for Aβ42 and $K_v$7.2 nanoclusters (violin plot of density, with a dashed line at the median and solid lines at the quartiles). The mean of Aβ42 puncta is significantly larger than that of Kv7.2, on average (two-tailed paired t-test, $p = 0.0001$, $t=4.116$, $df=54$). **(c)** Scatter plot of the number of $K_v$7.2 puncta vs. the volume of the largest Aβ42 puncta in the cropped ROI, showing no significant correlation between the two (simple linear regression, 95% CI of slope [-0.003940, 0.005947], $p = 0.6855$, $R^2 = 0.003118$). (**d)** Scatter plot of the mean $K_v$7.2 puncta volume vs. the mean Aβ42 puncta volume, illustrating a sublinear relationship but significant positive correlation between the two (simple linear regression, $R^2 = 0.2977$, $p <0.0001$, 95% CI of slope [0.09878, 0.2437]). Compare to the line y = x (blue). **(e)** Scatter plot of the non-overlapped volume for Aβ42 (gray squares) and $K_v7.2$ (black dots) vs. the mutually overlapped volume of the two proteins, compared to the line y = x (blue). The volume of $K_v7.2$ outside of Aβ42 is on average larger than the volume of Aβ42 outside of $K_v7.2$ (two-tailed t-test on the difference between the ratio of non-overlapped volume to overlapped volume for $K_v7.2$ and Aβ42, $p < 0.0001$). While the non-overlapped volume of Aβ42 is correlated to the mutually overlapped volume (95% CI of slope [0.07733, 0.1700], $R^2 = 0.3510$, $p<0.0001$), the volume of $K_v7.2$ outside of Aβ42 is not (95% CI of slope [0.03205, 0.5237], $R^2 = 0.05608$, $p = 0.0817$).

**Table. S1.** Numbers of technical replicates (synapses) used in Figure 2 analysis for each synaptic protein, layer, and mouse.

| **Protein** | **Layer** | **Animal** | **Number of Synapses** |
|---|---|---|---|
| Cav2.1 | 1 | A | 54 |
| | 2/3 | A | 55 |
| | 4 | A | 50 |
| | 1 | B | 51 |
| | 2/3 | B | 56 |
| | 4 | B | 52 |
| Bassoon | 1 | A | 55 |
| | 2/3 | A | 55 |
| | 4 | A | 55 |
| | 1 | B | 54 |
| | 2/3 | B | 55 |
| | 4 | B | 55 |
| Homer1 | 1 | A | 50 |

| | | | |
|---|---|---|---|
| | 2/3 | A | 55 |
| | 4 | A | 50 |
| | 1 | B | 50 |
| | 2/3 | B | 49 |
| | 4 | B | 50 |
| PSD95 | 1 | A | 49 |
| | 2/3 | A | 52 |
| | 4 | A | 60 |
| | 1 | B | 70 |
| | 2/3 | B | 55 |
| | 4 | B | 54 |
| RIM1/2 | 1 | A | 55 |
| | 2/3 | A | 52 |
| | 4 | A | 55 |
| | 1 | B | 55 |
| | 2/3 | B | 58 |
| | 4 | B | 55 |
| Shank3 | 1 | A | 49 |
| | 2/3 | A | 50 |
| | 4 | A | 44 |
| | 1 | B | 55 |
| | 2/3 | B | 50 |
| | 4 | B | 55 |
| SynGAP | 1 | A | 51 |
| | 2/3 | A | 50 |
| | 4 | A | 50 |

| | | | |
|---|---|---|---|
| | 1 | B | 49 |
| | 2/3 | B | 55 |
| | 4 | B | 50 |

**Table. S2.** Statistical analysis for Figure 2I: synapse decrowding analysis statistics of mean signal amplitude within dilated reference channel ROIs.

3-way analysis of variance (ANOVA) was used to test the statistical significance of the effect of multiple factors, including protein, pre- or post-expansion staining, and inside or outside of dilated reference ROIs as categorical variables, on the mean value of signal intensity inside ROIs. Further analysis was then conducted on selected significant factors. To determine which of the seven proteins had significantly different mean signal amplitude between pre- and post-expansion staining, multiple 2-way ANOVAs were run using linear models with inside/outside reference ROIs and pre/post staining as categorical variables. ANOVA p-values were corrected using the Holm-Sidak method. To compare the mean signal amplitude within dilated reference ROIs of pre- and post-expansion stained synapses, Sidak's multiple comparisons test, following one-way ANOVA, was run to compare the means of each pair of groups (pre/inside, pre/outside, post/inside, post/outside) in GraphPad Prism, only on those proteins that showed a significant effect of the pre/post expansion staining factor in 2-way ANOVA. For all ANOVA testing, the Python *statsmodels* package was used to run ordinary least squares models and calculate ANOVA tables with Type III sum of squares.

i) 3-way ANOVA table

| **Factor** | **SSE** | **df** | **F** | **Pr(>F)** |
|---|---|---|---|---|
| Intercept | 996713 | 1 | 13.2508 | 0.0004 |
| Protein | 3729370 | 6 | 8.2634 | 0.0000 |
| In/Out | 1855708 | 1 | 24.6708 | 0.0000 |
| Pre/Post | 1016339 | 1 | 13.5118 | 0.0003 |
| Residual | 11959805 | 159 | | |

ii) Multiple 2-way ANOVAs, with Holm-Sidak corrected p-values

| **Protein** | **Uncorrected: P(>F), Pre/Post Factor** | **Corrected: P(>F), Pre/Post Factor** | **Reject null?** | **Uncorrected: P(>F), Inside/Outside Factor** | **Corrected: P(>F), Inside/Outside Factor** | **Reject null?** |
|---|---|---|---|---|---|---|
| Bassoon | 0.0000 | 0.0001 | True | 0.0000 | 0.0000 | True |
| $Ca_v$2.1 | 0.0400 | 0.1152 | False | 0.0571 | 0.1110 | False |
| Homer1 | 0.2211 | 0.2211 | False | 0.0000 | 0.0001 | True |
| PSD95 | 0.0208 | 0.0806 | False | 0.1972 | 0.1972 | False |
| RIM1/2 | 0.0492 | 0.1152 | False | 0.0000 | 0.0000 | True |
| Shank3 | 0.0047 | 0.0235 | True | 0.0000 | 0.0000 | True |
| SynGAP | 0.0005 | 0.0030 | True | 0.0016 | 0.0047 | True |

iii) Sidak's multiple comparisons test following ordinary one-way ANOVA

| **Protein** | **Pre-Post Inside, Pr(>t)** | **Pre-Post Outside, Pr(>t)** | **Pre Inside-Outside, Pr(>t)** | **Post Inside-Outside, Pr(>t)** |
|---|---|---|---|---|
| Bassoon | <0.0001 | 0.0549 | 0.0010 | <0.0001 |
| $Ca_v$2.1 | 0.0415 | 0.9830 | 0.9954 | 0.0543 |
| Homer1 | 0.6019 | 0.9582 | 0.0019 | 0.0104 |
| PSD95 | 0.0538 | 0.8335 | >0.9999 | 0.2513 |
| RIM1/2 | 0.0053 | 0.9816 | 0.0627 | <0.0001 |
| Shank3 | 0.0004 | 0.9748 | <0.0001 | <0.0001 |
| SynGAP | <0.0001 | 0.8141 | 0.9750 | <0.0001 |

**Table. S3.** Statistical analysis for Figure 2J: synapse decrowding analysis of mean puncta volume within dilated reference channel ROIs.

3-way analysis of variance (ANOVA) was used to test the statistical significance of the effect of multiple factors, including protein, pre- or post-expansion staining, and inside or outside of dilated reference ROIs as categorical variables, on the mean value of ROI volume. Further analysis was then conducted on selected significant factors. To determine which of the seven proteins had significantly different mean ROI volume between pre- and post-expansion staining, multiple 2-way ANOVAs were run using linear models with inside/outside reference ROIs and pre/post staining as categorical variables. ANOVA p-values were corrected using the Holm-Sidak method. To compare the mean signal amplitude within dilated reference ROIs of pre- and post-expansion stained synapses, Sidak's multiple comparisons test, following one-way ANOVA, was run to compare the means of each pair of groups (pre/inside, pre/outside, post/inside, post/outside) in GraphPad Prism, only on those proteins that showed a significant effect of the pre/post expansion staining factor in 2-way ANOVA. For all ANOVA testing, the Python *statsmodels* package was used to run ordinary least squares models and calculate ANOVA tables with Type III sum of squares.

i) 3-way ANOVA table

| **Factor** | **SSE** | **df** | **F** | **Pr(>F)** |
|---|---|---|---|---|
| Intercept | 10412084 | 1 | 95.9970 | <0.0001 |
| Protein | 12105567 | 6 | 18.6017 | <0.0001 |
| In/Out | 20426451 | 1 | 188.3271 | <0.0001 |
| Pre/Post | 3357922 | 1 | 30.9593 | <0.0001 |
| Residual | 17245560 | 159 | | |

ii) Multiple 2-way ANOVAs, with Holm-Sidak corrected p-values

| **Protein** | **Uncorrected: P(>F), Pre/Post Factor** | **Corrected: P(>F), Pre/Post Factor** | **Reject null?** | **Uncorrected: P(>F), Inside/Outside Factor** | **Corrected: P(>F), Inside/Outside Factor** | **Reject null?** |
|---|---|---|---|---|---|---|
| Bassoon | <0.0001 | 0.0003 | True | <0.0001 | <0.0001 | True |
| $Ca_V2.1$ | 0.0025 | 0.0100 | True | 0.0889 | 0.1396 | False |
| Homer1 | 0.7501 | 0.7501 | False | <0.0001 | <0.0001 | True |

| | | | | | | |
|---|---|---|---|---|---|---|
| PSD95 | 0.0398 | 0.1148 | False | 0.0724 | 0.1396 | False |
| RIM1/2 | 0.0004 | 0.0021 | True | <0.0001 | <0.0001 | True |
| Shank3 | 0.3745 | 0.6087 | False | <0.0001 | <0.0001 | True |
| SynGAP | 0.0000 | 0.0002 | True | <0.0001 | <0.0001 | True |

iii) Sidak's multiple comparisons test following ordinary one-way ANOVA

| **Protein** | **Pre-Post Inside, Pr(>t)** | **Pre-Post Outside, Pr(>t)** | **Pre Inside-Outside, Pr(>t)** | **Post Inside-Outside, Pr(>t)** |
|---|---|---|---|---|
| Bassoon | <0.0001 | 0.3482 | 0.0007 | <0.0001 |
| $Ca_V$2.1 | 0.0010 | 0.8045 | 0.9755 | 0.0188 |
| Homer1 | 0.9210 | 0.9978 | <0.0001 | <0.0001 |
| PSD95 | 0.0250 | 0.9969 | >0.9999 | 0.0405 |
| RIM1/2 | <0.0001 | 0.7364 | 0.2598 | <0.0001 |
| Shank3 | 0.6744 | 0.9991 | <0.0001 | <0.0001 |
| SynGAP | <0.0001 | 0.2276 | 0.0001 | <0.0001 |

**Table. S4.** Accompanying statistics for Figure 2M-O and Fig. S5: analysis of distortion caused by post-expansion staining.

(i) Descriptive statistics for the number of puncta in the pre-expansion staining channel for Homer1 and Shank3 (Fig. S5d).

| | Homer1 | Shank3 |
|---|---|---|
| Number of values | 304 | 309 |
| Minimum | 0.000 | 0.000 |
| Maximum | 3.000 | 4.000 |

| | | |
|---|---|---|
| Range | 3.000 | 4.000 |
| Mean | 1.250 | 1.298 |
| Std. Deviation | 0.5296 | 0.5772 |
| Std. Error of Mean | 0.03038 | 0.03283 |
| Lower 95% CI of mean | 1.190 | 1.233 |
| Upper 95% CI of mean | 1.310 | 1.362 |

(ii) Descriptive statistics for the number of puncta in the post-expansion staining channel for Homer1 and Shank3 (Fig. S5e).

| | Homer1 | Shank3 |
|---|---|---|
| Number of values | 304 | 309 |
| Minimum | 1.000 | 0.000 |
| Maximum | 3.000 | 5.000 |
| Range | 2.000 | 5.000 |
| Mean | 1.174 | 1.285 |
| Std. Deviation | 0.4212 | 0.6214 |
| Std. Error of Mean | 0.02416 | 0.03535 |
| Lower 95% CI of mean | 1.127 | 1.215 |
| Upper 95% CI of mean | 1.222 | 1.354 |

(iii) Descriptive statistics for the difference in the number of puncta in the post-expansion staining channels for Homer1 and Shank3 (Fig. S5f).

| | Homer1 | Shank3 |
|---|---|---|
| Number of values | 304 | 309 |
| Minimum | -2.000 | -2.000 |

| | | |
|---|---|---|
| Maximum | 2.000 | 3.000 |
| Range | 4.000 | 5.000 |
| Mean | -0.07566 | -0.01294 |
| Std. Deviation | 0.5838 | 0.6292 |
| Std. Error of Mean | 0.03348 | 0.03580 |
| Lower 95% CI of mean | -0.1415 | -0.08338 |
| Upper 95% CI of mean | -0.009772 | 0.05749 |

(iv) Descriptive statistics for the difference in the number of puncta in the post-expansion staining channels, normalized to the number of puncta in the pre-expansion staining channel, for Homer1 and Shank3 (Fig. 2M). Values with zero puncta in the pre-expansion staining channel were excluded.

| | Homer1 | Shank3 |
|---|---|---|
| Number of values | 303 | 306 |
| Minimum | -0.6667 | -1.000 |
| Maximum | 2.000 | 3.000 |
| Range | 2.667 | 4.000 |
| Mean | 0.01540 | 0.04630 |
| Std. Deviation | 0.3756 | 0.4348 |
| Std. Error of Mean | 0.02158 | 0.02485 |
| Lower 95% CI of mean | -0.02706 | -0.002610 |
| Upper 95% CI of mean | 0.05786 | 0.09520 |

(v) Descriptive statistics for the difference in half-maximum correlation shift between pre-post pixel-wise correlation and pre-pre pixel-wise autocorrelation (Fig. 2N).

| | Homer1 | Shank3 |
|---|---|---|

| | | |
|---|---|---|
| Number of values | 909 | 915 |
| Minimum | -100.0 | -100.0 |
| Maximum | 64.90 | 65.50 |
| Range | 164.9 | 165.5 |
| Mean | 0.3061 | 5.986 |
| Std. Deviation | 14.84 | 16.15 |
| Std. Error of Mean | 0.4923 | 0.5340 |
| Lower 95% CI of mean | -0.6601 | 4.938 |
| Upper 95% CI of mean | 1.272 | 7.034 |

(vi) Descriptive statistics for the difference in half-maximum correlation shift between pre-post pixel-wise correlation and post-post pixel-wise autocorrelation (Fig. 2O).

| | Homer1 | Shank3 |
|---|---|---|
| Number of values | 909 | 915 |
| Minimum | -100.0 | -100.0 |
| Maximum | 44.60 | 68.50 |
| Range | 144.6 | 168.5 |
| Mean | 4.523 | 6.607 |
| Std. Deviation | 17.07 | 16.03 |
| Std. Error of Mean | 0.5662 | 0.5298 |
| Lower 95% CI of mean | 3.412 | 5.567 |
| Upper 95% CI of mean | 5.634 | 7.647 |

**Table. S5.** Statistical analysis for Figure 2k: synaptic puncta analysis for mean volume of the largest synaptic connected components.

3-way analysis of variance (ANOVA) was used to test the statistical significance of the effect of multiple factors, including protein, layer, and pre- or post-expansion staining as categorical variables, on the mean value of synapse volume. Categorical variables for interactions between protein and pre/post staining and layer and protein were included in the linear model based on biological feasibility of these interactions. Further analysis was then conducted on selected significant factors. To determine which of the seven proteins had significantly different volume between pre- and post-expansion staining, multiple 2-way ANOVAs were run using linear models with layer and pre/post staining as categorical variables. ANOVA p-values were corrected using the Holm-Sidak method. To compare the volume of pre- and post-expansion stained synapses for each of the seven proteins, three t-tests (one for each layer) were run (see **Table. S1** for numbers of biological and technical replicates). Statistical significance was determined using the Holm-Sidak method, with alpha = 0.05. Each layer was analyzed individually, without assuming a consistent standard deviation. For all ANOVA testing, the Python *statsmodels* package was used to run ordinary least squares models and calculate ANOVA tables with Type III sum of squares.

i) 3-way ANOVA table

| **Factor** | **SSE** | **df** | **F** | **Pr(>F)** |
|---|---|---|---|---|
| Intercept | 1692606 | 1 | 47.0052 | <0.0001 |
| Protein | 6209794 | 6 | 28.7420 | 0.0000 |
| Layer | 239338 | 2 | 3.3233 | 0.0433 |
| Pre/Post | 1287591 | 1 | 35.7576 | <0.0001 |
| Protein-Pre/Post Interaction | 2846212 | 6 | 13.1737 | <0.0001 |
| Layer/Protein Interaction | 710693 | 12 | 1.6447 | 0.1055 |
| Residual | 2016497 | 56 | | |

ii) Multiple 2-way ANOVAs, with Holm-Sidak corrected p-values

| **Protein** | **Uncorrected: P(>F), Pre/Post Factor** | **Corrected: P(>F), Pre/Post Factor** | **Reject null?** |
|---|---|---|---|

| | | | |
|---|---|---|---|
| Bassoon | <0.0001 | <0.0001 | True |
| Cav2.1 | 0.0004 | 0.0018 | True |
| Homer1 | 0.9497 | 0.9497 | False |
| PSD95 | 0.0183 | 0.0540 | False |
| RIM1/2 | <0.0001 | 0.0002 | True |
| Shank3 | 0.4284 | 0.6733 | False |
| SynGAP | <0.0001 | 0.0001 | True |

iii) Multiple t-tests, with Holm-Sidak corrected p-values

| **Protein** | **Layer** | **Mean Difference** | **SE of difference** | **t ratio** | **df** | **Adjusted p value** |
|---|---|---|---|---|---|---|
| Bassoon | L1 | -932.5 | 144.7 | 6.443 | 107.0 | <0.000001 |
| | L2/3 | -1040 | 147.6 | 7.045 | 108.0 | <0.000001 |
| | L4 | -1082 | 148.8 | 7.272 | 108.0 | <0.000001 |
| Cav2.1 | L1 | -424.0 | 58.19 | 7.287 | 103.0 | <0.000001 |
| | L2/3 | -588.0 | 61.76 | 9.521 | 109.0 | <0.000001 |
| | L4 | -953.4 | 88.30 | 10.80 | 103.0 | <0.000001 |
| Homer1 | L1 | 37.17 | 162.8 | 0.2283 | 98.00 | 0.819888 |
| | L2/3 | 166.3 | 140.9 | 1.181 | 102.0 | 0.540672 |
| | L4 | -181.5 | 149.8 | 1.212 | 101.0 | 0.540672 |
| PSD95 | L1 | -306.8 | 28.19 | 10.89 | 117.0 | <0.000001 |
| | L2/3 | -543.2 | 35.78 | 15.18 | 106.0 | <0.000001 |
| | L4 | -586.4 | 41.31 | 14.19 | 112.0 | <0.000001 |
| RIM1/2 | L1 | -451.4 | 102.9 | 4.389 | 108.0 | 0.000027 |
| | L2/3 | -696.1 | 103.0 | 6.758 | 111.0 | <0.000001 |
| | L4 | -632.9 | 94.00 | 6.733 | 108.0 | <0.000001 |

| | | | | | | |
|---|---|---|---|---|---|---|
| Shank3 | L1 | -55.09 | 159.0 | 0.3465 | 97.00 | 0.926942 |
| | L2/3 | -42.60 | 175.5 | 0.2427 | 98.00 | 0.926942 |
| | L4 | -127.9 | 130.8 | 0.9781 | 108.0 | 0.699554 |
| SynGAP | L1 | -644.2 | 136.0 | 4.736 | 97.00 | 0.000007 |
| | L2/3 | -1032 | 138.1 | 7.470 | 103.0 | <0.000001 |
| | L4 | -1233 | 100.8 | 12.24 | 98.00 | <0.000001 |

**Table. S6.** Statistical analysis for Figure 2L: synaptic puncta analysis of mean signal-to-noise ratio within the largest synaptic connected components.

3-way analysis of variance (ANOVA) was used to test the statistical significance of the effect of multiple factors, including protein, layer, and pre- or post-expansion staining as categorical variables, on the mean value of synapse SNR. Categorical variables for interactions between protein and pre/post staining and layer and protein were included in the linear model based on biological feasibility of these interactions. Further analysis was then conducted on selected significant factors. To determine which of the seven proteins had significantly different SNR between pre- and post-expansion staining, multiple 2-way ANOVAs were run using linear models with layer and pre/post staining as categorical variables. ANOVA p-values were corrected using the Holm-Sidak method. To compare the SNR of pre- and post-expansion stained synapses for each of the seven proteins, three t-tests (one for each layer) were run (n = 50 puncta per layer, n = 2 mice). Statistical significance was determined using the Holm-Sidak method, with alpha = 0.05. Each layer was analyzed individually, without assuming a consistent standard deviation. For all ANOVA testing, the Python *statsmodels* package was used to run ordinary least squares models and calculate ANOVA tables with Type III sum of squares.

i) 3-way ANOVA table

| **Factor** | **SSE** | **df** | **F** | **PR(>F)** |
|---|---|---|---|---|
| Intercept | 661.26 | 1 | 217.76 | <0.0001 |
| Protein | 188.08 | 6 | 10.322 | <0.0001 |
| Layer | 20.11 | 2 | 3.311 | 0.0438 |
| Pre/Post | 60.84 | 1 | 20.033 | <0.0001 |

| | | | | |
|---|---|---|---|---|
| Protein-Pre/Post Interaction | 203.51 | 6 | 11.170 | <0.0001 |
| Layer/Protein Interaction | 29.21 | 12 | 0.802 | 0.6472 |
| Residual | 170.06 | 56 | | |

ii) Multiple 2-way ANOVAs, with Holm-Sidak corrected p-values

| **Protein** | **Uncorrected: P(>F), Pre/Post Factor** | **Corrected: P(>F), Pre/Post Factor** | **Reject null?** |
|---|---|---|---|
| Bassoon | <0.0001 | <0.0001 | True |
| Cav2.1 | 0.0017 | 0.0052 | True |
| Homer1 | 0.0101 | 0.0201 | True |
| PSD95 | 0.0001 | 0.0005 | True |
| RIM1/2 | 0.0000 | 0.0001 | True |
| Shank3 | 0.1281 | 0.1281 | False |
| SynGAP | <0.0001 | <0.0001 | True |

iii) Multiple t-tests, with Holm-Sidak corrected p-values

| **Protein** | **Layer** | **Mean Difference** | **SE of difference** | **t ratio** | **df** | **Adjusted p value** |
|---|---|---|---|---|---|---|
| Bassoon | L1 | -5.510 | 0.4489 | 12.27 | 107.0 | <0.000001 |
| | L2/3 | -5.568 | 0.4563 | 12.20 | 108.0 | <0.000001 |
| | L4 | -6.261 | 0.4519 | 13.85 | 108.0 | <0.000001 |
| $Ca_V2.1$ | L1 | -3.153 | 0.4835 | 6.522 | 103.0 | <0.000001 |
| | L2/3 | -4.168 | 0.4344 | 9.594 | 109.0 | <0.000001 |
| | L4 | -6.189 | 0.5956 | 10.39 | 103.0 | <0.000001 |

| | | | | | | |
|---|---|---|---|---|---|---|
| Homer1 | L1 | -4.328 | 0.7455 | 5.805 | 98.00 | <0.000001 |
| | L2/3 | -4.194 | 0.5870 | 7.145 | 102.0 | <0.000001 |
| | L4 | -5.680 | 0.5969 | 9.516 | 101.0 | <0.000001 |
| PSD95 | L1 | -4.219 | 0.4373 | 9.649 | 117.0 | <0.000001 |
| | L2/3 | -6.052 | 0.7300 | 8.290 | 106.0 | <0.000001 |
| | L4 | -6.041 | 0.5395 | 11.20 | 112.0 | <0.000001 |
| RIM1/2 | L1 | -4.435 | 0.4005 | 11.07 | 108.0 | <0.000001 |
| | L2/3 | -3.962 | 0.4678 | 8.468 | 111.0 | <0.000001 |
| | L4 | -3.179 | 0.5557 | 5.721 | 108.0 | <0.000001 |
| Shank3 | L1 | -3.417 | 0.7348 | 4.650 | 97.00 | 0.000031 |
| | L2/3 | -2.151 | 0.6277 | 3.427 | 98.00 | 0.001784 |
| | L4 | -2.002 | 0.6369 | 3.143 | 108.0 | 0.002162 |
| SynGAP | L1 | -10.67 | 0.7954 | 13.42 | 97.00 | <0.000001 |
| | L2/3 | -12.22 | 0.8446 | 14.47 | 103.0 | <0.000001 |
| | L4 | -15.82 | 0.9348 | 16.93 | 98.00 | <0.000001 |

**Table. S7.** Statistical analysis for Fig. S2: comparison of the effects of antigen retrieval and ExR procedures on mean signal intensity within and outside of synaptic puncta.

To compare the mean signal amplitude within dilated reference ROIs of antigen retrieval- and non antigen retrieval-treated synapses, Sidak's multiple comparisons test was run to compare the means of samples treated with ExR and samples treated with antigen retrieval followed by pre-expansion staining (AR) in the foreground or background, as defined by the reference channel, in GraphPad Prism. 2-way ANOVA was run in GraphPad Prism using Type III sum of squares.

i) 2-way ANOVA table for $Ca_v2.1$.

| Factor | SSE | DF | F (DFn, DFd) | Pr(>F) |
|---|---:|---:|---:|---:|
| Interaction | 4380 | 3 | F (3, 232) = 35.20 | P<0.0001 |

| | | | | |
|---|---|---|---|---|
| Antigen retrieval | 2939 | 1 | F (1, 232) = 70.85 | P<0.0001 |
| Pre/post-expansion staining | 84306 | 3 | F (3, 232) = 677.6 | P<0.0001 |
| Residual | 9622 | 232 | | |

ii) Sidak's multiple comparisons test on the difference between ExR (decrowding)  and antigen retrieval followed by pre-expansion staining (AR) for $Ca_v$2.1, grouped by location inside dilated reference ROIs (foreground) or outside dilated reference ROIs (background).

| **Group** | **Mean Diff.** | **95.00% CI of diff.** | **Significant?** | **Adjusted P Value** |
|---|---|---|---|---|
| Foreground | 35.40 | 30.16 to 40.64 | Yes | <0.0001 |
| Background | 13.66 | 8.420 to 18.91 | Yes | <0.0001 |

iii) 2-way ANOVA table for Homer1.

| **Factor** | **SSE** | **DF** | **F (DFn, DFd)** | **Pr(>F)** |
|---|---|---|---|---|
| Interaction | 98791 | 3 | F (3, 232) = 328.4 | P<0.0001 |
| Antigen retrieval | 60058 | 1 | F (1, 232) = 598.9 | P<0.0001 |
| Pre/post-expansion staining | 2437902 | 3 | F (3, 232) = 8104 | P<0.0001 |
| Residual | 23264 | 232 | | |

iv) Sidak's multiple comparisons test on the difference between decrowding (ExR) and antigen retrieval followed by pre-expansion staining (AR) for Homer1, grouped by location inside dilated reference ROIs (foreground) or outside dilated reference ROIs (background).

| **Group** | **Mean Diff.** | **95.00% CI of diff.** | **Significant?** | **Adjusted P Value** |
|---|---|---|---|---|
| Foreground | -142.3 | -150.5 to -134.2 | Yes | <0.0001 |
| Background | -26.61 | -34.77 to -18.46 | Yes | <0.0001 |

v) 2-way ANOVA table for PSD95.

| **Factor** | **SSE** | **DF** | **F (DFn, DFd)** | **Pr(>F)** |
|---|---|---|---|---|
| Interaction | 113634 | 3 | F (3, 232) = 809.4 | P<0.0001 |
| Antigen retrieval | 45778 | 1 | F (1, 232) = 978.2 | P<0.0001 |
| Pre/post-expansion staining | 2007684 | 3 | F (3, 232) = 14301 | P<0.0001 |
| Residual | 10857 | 232 | | |

vi) Sidak's multiple comparisons test on the difference between decrowding (ExR) and antigen retrieval followed by pre-expansion staining (AR) for PSD95, grouped by location inside dilated reference ROIs (foreground) or outside dilated reference ROIs (background).

| **Group** | **Mean Diff.** | **95.00% CI of diff.** | **Significant?** | **Adjusted P Value** |
|---|---|---|---|---|
| Foreground | 271.5 | 265.9 to 277.1 | Yes | <0.0001 |
| Background | 37.87 | 32.31 to 43.44 | Yes | <0.0001 |

**Table. S8.** Statistical analysis for Fig. S2: comparison of the effects of antigen retrieval and ExR procedures on total signal volume within and outside of synaptic puncta.

To compare the total volume within dilated reference ROIs of antigen retrieval- and non antigen retrieval-treated synapses, Sidak's multiple comparisons test was run to compare the means samples treated with ExR and samples treated with antigen retrieval followed by pre-expansion staining (AR) in the foreground or background, as defined by the reference channel, in GraphPad Prism. 2-way ANOVA was run in GraphPad Prism using Type III sum of squares.

i) 2-way ANOVA table for $Ca_v$2.1.

| **Factor** | **SSE** | **DF** | **F (DFn, DFd)** | **Pr(>F)** |
|---|---|---|---|---|
| Interaction | 3664 | 3 | F (3, 232) = 0.5364 | P=0.6578 |
| Antigen retrieval | 3496 | 1 | F (1, 232) = 1.536 | P=0.2165 |

| | | | | |
|---|---|---|---|---|
| Pre/post-expansion staining | 9616433 | 3 | F (3, 232) = 1408 | P<0.0001 |
| Residual | 528212 | 232 | | |

ii) Sidak's multiple comparisons test on the difference between decrowding (ExR) and antigen retrieval followed by pre-expansion staining (AR) for $Ca_v2.1$, grouped by location inside dilated reference ROIs (foreground) or outside dilated reference ROIs (background).

| **Group** | **Mean Diff.** | **95.00% CI of diff.** | **Significant?** | **Adjusted P Value** |
|---|---|---|---|---|
| Foreground | 458.9 | 420.1 to 497.7 | Yes | <0.0001 |
| Background | 0.6667 | -38.18 to 39.51 | No | >0.9999 |

iii) 2-way ANOVA table for Homer1.

| **Factor** | **SSE** | **DF** | **F (DFn, DFd)** | **Pr(>F)** |
|---|---|---|---|---|
| Interaction | 9272076 | 3 | F (3, 232) = 68.37 | P<0.0001 |
| Antigen retrieval | 2924938 | 1 | F (1, 232) = 64.71 | P<0.0001 |
| Pre/post-expansion staining | 635644935 | 3 | F (3, 232) = 4687 | P<0.0001 |
| Residual | 10487186 | 232 | | |

iv) Sidak's multiple comparisons test on the difference between decrowding (ExR) and antigen retrieval followed by pre-expansion staining (AR) for Homer1, grouped by location inside dilated reference ROIs (foreground) or outside dilated reference ROIs (background).

| **Group** | **Mean Diff.** | **95.00% CI of diff.** | **Significant?** | **Adjusted P Value** |
|---|---|---|---|---|
| Foreground | 285.7 | 112.6 to 458.8 | Yes | <0.0001 |
| Background | -124.6 | -297.7 to 48.48 | No | 0.4955 |

v) 2-way ANOVA table for PSD95.

| Factor | SSE | DF | F (DFn, DFd) | Pr(>F) |
|---|---|---|---|---|
| Interaction | 18508762 | 3 | F (3, 232) = 266.9 | P<0.0001 |
| Antigen retrieval | 8905439 | 1 | F (1, 232) = 385.3 | P<0.0001 |
| Pre/post-expansion staining | 667591009 | 3 | F (3, 232) = 9628 | P<0.0001 |
| Residual | 5362087 | 232 | | |

vi) Sidak's multiple comparisons test on the difference between decrowding (ExR) and antigen retrieval followed by pre-expansion staining (AR) for PSD95, grouped by location inside dilated reference ROIs (foreground) or outside dilated reference ROIs (background).

| Group | Mean Diff. | 95.00% CI of diff. | Significant? | Adjusted P Value |
|---|---|---|---|---|
| Foreground | 4676 | 4552 to 4800 | Yes | <0.0001 |
| Background | 690.8 | 567.0 to 814.6 | Yes | <0.0001 |
